# Cyclic peptide inhibitors stabilize Gq/11 heterotrimers

**DOI:** 10.1101/2023.10.24.563737

**Authors:** Jonas Mühle, Matthew J. Rodrigues, Judith Alenfelder, Lars Jürgenliemke, Ramon Guixà-González, Arianna Bacchin, Fabio Andres, Michael Hennig, Hannes Schihada, Max Crüsemann, Gabriele M. König, Evi Kostenis, Gebhard Schertler, Xavier Deupi

## Abstract

Heterotrimeric G proteins play a central role in cellular signaling, acting as switchable molecular regulators. Consequently, pharmacological agents to control G protein activity are of utmost importance to advance our understanding of this signal transduction system. The natural depsipeptides FR900359 (FR) and YM-254890 (YM) are two highly specific and widely used inhibitors of heterotrimeric Gq/11 proteins. These compounds have traditionally been understood to inhibit GDP dissociation by preventing the separation of the GTPase and α-helical domains of the Gα subunit. In this work, we have determined the high-resolution crystal structures of FR and YM bound to heterotrimeric G11 and used them to explain the molecular basis underlying their efficient suppression of G protein signaling. Notably, our data show that FR and YM also function as stabilizers of the interface between the Gα and Gβ subunits, acting as ‘molecular adhesives’ that stabilize the entire heterotrimer. Our results reveal unrecognized mechanistic features that explain how FR and YM effectively blunt Gq/11 signaling in living cells.

## INTRODUCTION

A precise cellular signaling machinery is a prerequisite to maintaining homeostasis in living systems. Heterotrimeric guanine nucleotide-binding proteins (G proteins) play a crucial role in the G protein-coupled receptor (GPCR) signaling pathway. In this signal transduction process, a physicochemical signal from the environment is relayed into the cell through the activation of transmembrane GPCRs resulting in the recruitment and activation of intracellular downstream effectors. The primary effectors of GPCRs, G proteins, exist as heterotrimers composed of a GDP-bound Gα subunit bound tightly to a constitutive Gβγ dimer (“OFF state”). G proteins can be divided into four main classes based on sequence homology and the signaling profile of their alpha subunit: Gαi, Gαs, Gαq/11, and Gα12/13. Upon recruitment to an active receptor (or to a non-receptor guanine nucleotide exchange factor), conformational changes in Gα lead to nucleotide exchange from GDP to GTP resulting in the dissociation of the trimer into a GTP-bound Gα subunit and a Gβγ complex (“ON state”). Both subunits can then modulate downstream signaling partners until the Gα_GTP_ subunit hydrolyzes GTP back to GDP (assisted by regulator of G protein signaling (RGS) proteins), thus initiating the recovery of the heterotrimer and allowing additional cycles of G protein activation^1, 2, 3^.

While there is a plethora of natural and synthetic substances capable of modulating GPCR function, only a few are known for G proteins. The recent discovery of two small molecule-like Gαs inhibitors – GD20 and GN13– illustrates the interest in the development of subfamily-specific and state-selective G protein modulators as research tools as well as potential therapeutic agents^4^. The most potent known cell-permeable G protein inhibitors are the macrocyclic depsipeptide natural products FR900359 (FR) and YM-254890 (YM)^5^, which specifically inhibit GDP dissociation in the Gαq/11 family^6, 7, 8, 9^. Both depsipeptides are chemically very similar and differ only in two side chains: anchors 1 and 2 (**Fig. 1a**). FR has bulkier ethyl and isopropyl groups in anchors 1 and 2, respectively, instead of the two smaller methyl groups of YM. Numerous preclinical studies have corroborated the power of these inhibitors as pharmacological research tools to investigate G protein signaling as well as their potential for the treatment of several diseases including obesity, metabolic disorders, asthma, and cancer^10, 11, 12, 13, 14, 15^.

A 2.9 Å resolution crystal structure of YM bound to a heterotrimeric Gαq/i chimera provided initial insights into the mechanism of action of this inhibitor. YM binds at the interface between the RAS-like and the α-helical domains of Gα in a shallow binding pocket that includes the two interdomain junction sites “linker 1” and “linker 2/switch I”. Stabilization of this interface is suggested to cause the observed slower GDP-to-GTP exchange induced by YM^16^. Key residues in Gαq have been identified in the binding site, such as Arg60 (which rigidifies linker 1 through a hydrogen bond network) and Val184/Ile190 (which stabilize linker 2/switch I)^7^. This crystal structure has allowed protein engineering studies to create FR/YM-insensitive Gαq variants^17, 18^, as well as FR/YM-sensitive variants of other G protein families^11, 17, 19^. However, structure-based drug design efforts to create improved versions of FR and YM or to find new inhibitors selective for other G protein subfamilies have not been successful so far^20, 21, 22, 23^. There are other aspects of the FR/YM mechanism of action that are not yet well understood. For instance, YM has been shown to enhance Gβγ binding to Gαq in living cells through a yet unknown mechanism^24^, but whether Gβγ contributes directly to the action of YM or FR is elusive and cannot be concluded from the available crystal structure.

We postulated that the availability of a higher resolution structure of an FR-bound G protein would not only advance our mechanistic understanding of the mode of action of cyclic depsipeptide inhibitors, but also resolve any potential contribution of the Gβγ subunit complex. Thus, in this manuscript, we reveal two high-resolution X-ray structures of the Gα11β1γ2 heterotrimer bound to FR and YM. These constitute the first X-ray structures of a Gα11 protein. Both inhibitors adopt a very similar pose in the binding pocket, but slight structural differences in the G protein can be observed near the two anchor sites. Additionally, our high-resolution data enabled us to thoroughly refine all protein-ligand interactions and identify previously unknown contacts. Remarkably, our structures reveal a previously unidentified interaction of the inhibitors with Arg96 in the Gβ subunit. Thermostability and cellular BRET assays confirm that this interaction contributes to the stabilization of the G protein heterotrimer –and not only the Gα subunit. Thus, we propose that the stabilization of the Gα:Gβ interface is an essential part of the inhibition mechanism of FR and YM.

## RESULTS

We expressed and purified **(Suppl. Fig. 1)** different variants of Gαq/11 proteins as previously described^7^ (see Methods) to attempt the production of diffraction-quality crystals. We could successfully grow crystals of a heterotrimeric Gα11β1γ2 complex bound to FR using the vapor diffusion method at 4 °C and solve its structure by X-ray cryo-crystallography to a resolution of 1.43 Å **(Suppl. Table 1)**. The construct design was based on previously reported structures of the Gαq/11 family^7, 25^. Namely, the N-terminal helix of Gα11 was replaced by the first 29 amino acids of Gαi1 (Gα11iN1-29) to increase protein solubility. To optimize expression and purification, an N-terminal cleavable eGFP was fused to Gα11iN1-29. Gβ1 was N-terminally fused to a cleavable deca-histidine-tag. The prenylation site of Gγ2 was removed by introducing the known mutation C68S to create a fully soluble heterotrimer. The final crystallized complex was thus Gα11iN1-29β1γ2C68S:FR900359 which will be abbreviated from now on as G11iN_3_-S:FR (3=heterotrimer; S=soluble).

We assessed the functionality of our construct using several methods. First, we measured the thermostability of G11iN_3_-S in the apo state and the presence of FR (**Fig. 1b**). In the apo state, two transitions are observed in the melting temperature curve: one at 51.4 °C that corresponds to the unfolding of the Gɑ subunit and one at 57.5 °C that represents the unfolding of Gβγ. Upon FR binding, the unfolding curve simplifies to display a single transition with an inflection point at 60.6 °C. This significant increase in melting temperature in the presence of FR indicates specific ligand binding to functionally folded protein. Moreover, the change to a single melting point that is higher than the two individual melting temperatures in the apo state suggests the unpredicted stabilization of the Gα/Gβ interface –and, thus, of the entire heterotrimer– by FR. In addition, FR binding data to G11iN_3_-S measured by grating-coupled interferometry (GCI) shows nanomolar binding affinity under the assumption of a 1:1 binding mode (**Fig. 1c**), in close agreement with values reported in the literature^26^. Finally, we assessed the functionality of the purified protein by measuring its GTPase activity. The data show specific receptor-induced activation and FR-mediated inhibition of our construct (**Fig. 1d**).

**Figure 1:**
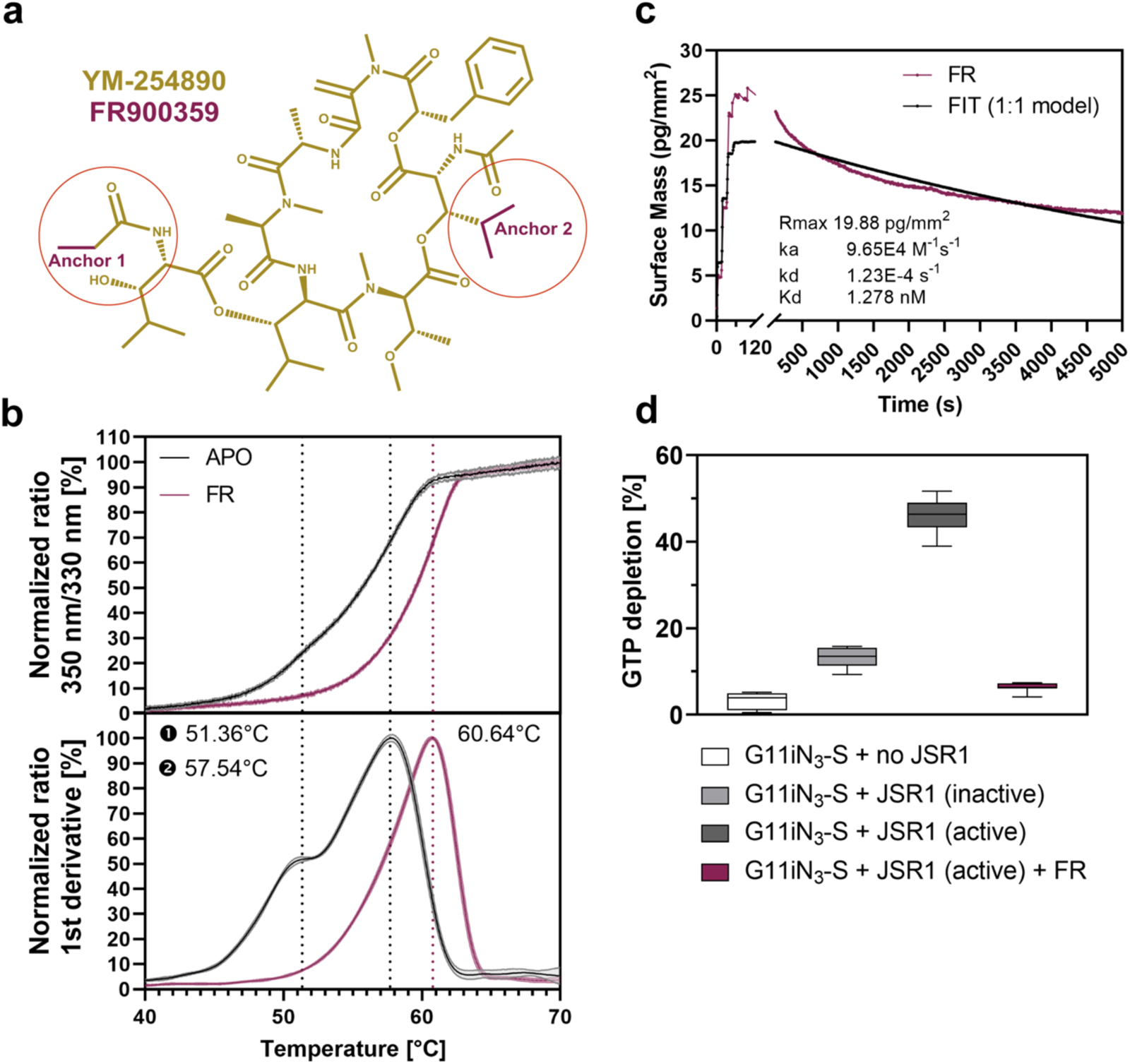
Ligand binding and functional characterization of the G11/FR complex. **a)** 2D representation of YM-254890 (YM) in dark yellow. FR900359 (FR) has the same scaffold as YM but with two extra moieties at anchors 1 and 2, shown and highlighted in red. Interestingly, the two anchors are located on opposite sides of the inhibitor scaffold. **b)** Thermostability assay of G11iN3-S in the apo state and in the presence of FR measuring the ratio of 350 nm and 330 nm tryptophan fluorescence. The upper panel shows the unfolding curves and the lower panel displays their first derivative, in which the peaks (maxima and saddle points) represent the melting temperature of each species. In the apo state, two transitions are observed: one at 51.36 ± 0.16 °C and one at 57.54 ± 0.11 °C. Upon FR binding, the unfolding curve changes to a single-transition curve with a point of inflection at 60.64± 0.18 *°C.* **c)** FR binding (120 s) and unbinding (5000 s) to G11iN3-S measured by grating-coupled interferometry (GCI) using the waveRAPID® technology. A single-digit nanomolar binding affinity was determined under the assumption of a 1:1 binding model with some uncertainty as the unbinding seems to be biphasic. However, it is unclear whether or not this is due to unspecific effects or part of the dissociation mechanism. **d)** GTPase Glo assay® measuring specific receptor-induced activation and FR-mediated inhibition of purified G11iN3-S. The protein shows some basal activity of GTP depletion (white) that increases moderately in the presence of an inactive GPCR (light grey). We used the light-activated GPCR jumping spider rhodopsin 1 (JSR1) to stimulate the G protein and observed a significant increase in GTP depletion rate (dark grey) that can be fully inhibited by the addition of FR (red).

Our model represents the first X-ray structure of a Gα11 protein as well as the first experimentally determined structure of a G protein bound to FR (**Fig. 2a**). The heterotrimeric G protein displays its typical fold, with the Gα subunit composed of a Ras-like GTPase and a α-helical domain connected by linker 1 and linker 2/switch I, a Gβ subunit with a seven-blade β-propeller fold and an α-helical N-terminus parallel to the Gγ subunit which further wraps around Gβ. The overall structure superposes well with other structures of the Gαq/11 family (e.g., 0.8 Å RMSD to the Gαq/i heterotrimer bound to YM^7^).

FR binds to a central part of the heterotrimer, at the interface of the two Gα domains and the Gβ subunit and near the GDP binding pocket, in the same site where YM-254890 binds to Gq/i^7^. The high-resolution diffraction data and the resulting detailed and well-defined electron density map **(Suppl. Fig. 2)** allowed the unambiguous modeling of the inhibitor in the binding pocket.

**Figure 2:**
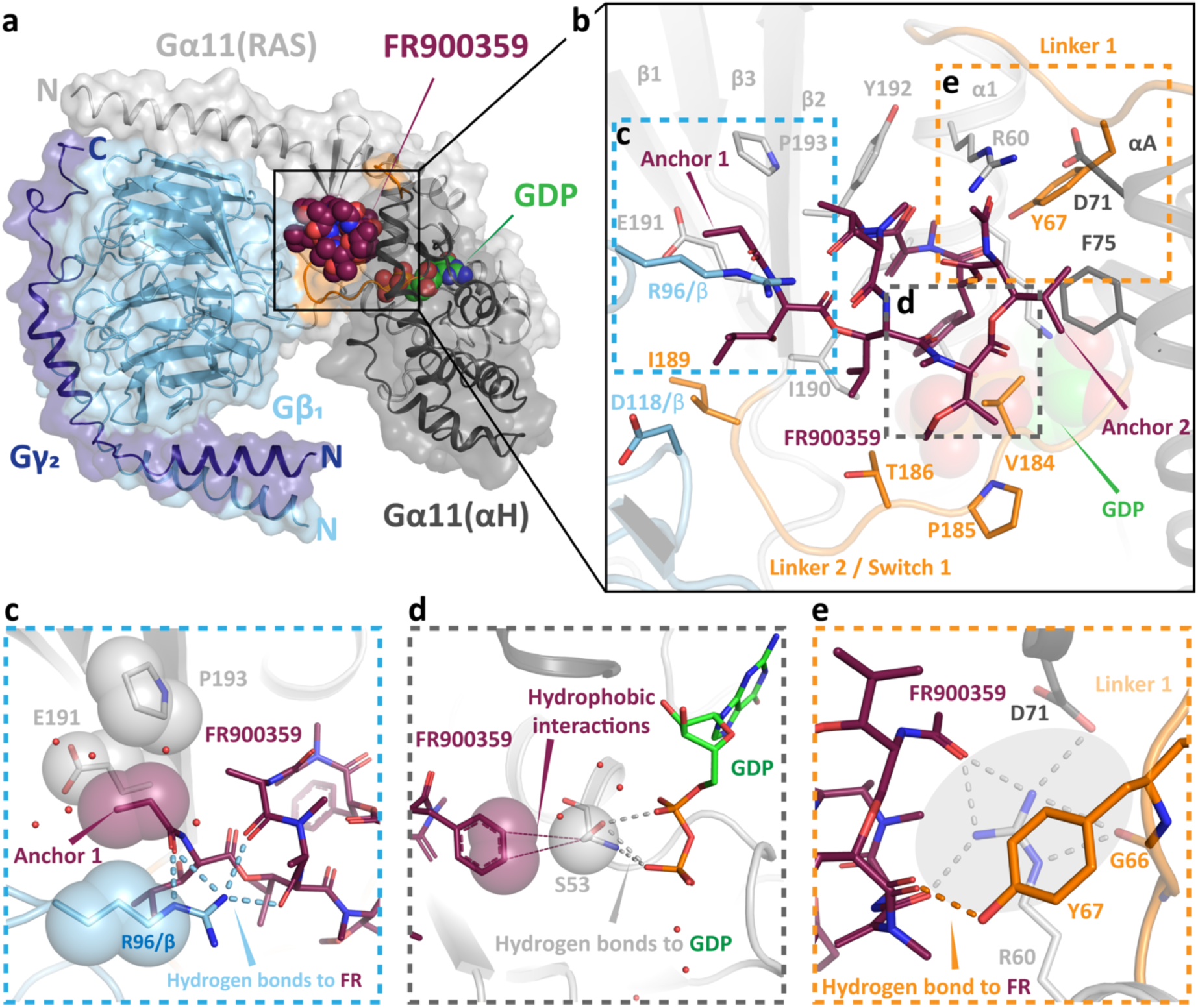
X-ray structure of G11iN3-S in complex with FR. **a)** Overall view of the crystal structure shown as cartoons with a translucent molecular surface. The Gα11 subunit is shown in grey (Gα-RAS: light; Gα-αH: dark), Gβ in cyan, Gγ in dark blue, and linker 1 and linker 2/switch I in orange. FR and GDP are depicted as spheres with red and green carbon atoms, respectively. **b)** Close-up view of the FR binding pocket. FR and the interacting protein side chains (distance < 3 Å) are depicted as sticks. The inhibitor interacts with residues from both Gα domains, with both linkers, and, additionally, contacts R96 from the Gβ subunit (blue dashed box). Two additional dashed boxes (grey, orange) highlight other remarkable ligand-protein interactions, which are shown in greater detail in panels c-e. **c)** The side chain of R96 in the Gβ subunit extends towards FR in a straight conformation and establishes four direct hydrogen bonds with the inhibitor (blue dashed lines). Furthermore, anchor 1 of FR inserts into a small hydrophobic pocket (shown as translucent spheres) composed of the R96 side chain in Gβ and E191 and P193 in Gɑ. **d)** Direct interaction of FR with the nucleotide-binding pocket: the phenyl ring of FR contacts S53 at the P-loop through a hydrophobic interaction (shown as red translucent spheres and thin lines). Residue S53 also coordinates the phosphate groups of GDP through hydrogen bonds (dashed lines). **e)** The direct polar contact between FR and Y67 in linker 1 (red dashed line) is shown in the foreground. The inhibitor also stabilizes linker 1 through a hydrogen bond network mediated by R60, shown in grey (circled dashed lines) in the background.

FR binds to a shallow pocket predominantly composed of helix α1 in the GTPase domain, helix αA in the α-helical domain, and of the joining linker 1 and linker 2/switch I. Furthermore, anchor 1 of FR extends into a wide-open cleft on the Gα/Gβ interface (**Fig. 2b and 2c**). Many putative interactions between FR and Gq have been predicted and tested^26^ and many of these are indeed observed in our structure. However, our high-resolution data allows us to detect new interactions that provide fresh data on the singular properties of FR.

The most striking feature in the binding pocket is a previously unidentified interaction of Arg96 in the Gβ subunit with the anchor 1 region of FR, consisting of four direct and one water-mediated hydrogen bonds (**Fig. 2c**). This interaction is responsible for four of the eleven polar contacts formed with the ligand and presumably contributes decisively to the stability of the heterotrimer. From a structural point of view, Arg96/Gβ appears to “cover” anchor 1 of FR locking it in a stable pose in an otherwise shallow binding pocket. The ethyl group of anchor 1 is also positioned near Glu191^G.S2.03^ (the superscript corresponds to the G protein general residue number^27^) and Pro193^G.S2.05^ of the GTPase domain. Hydrophobic interactions within this triad of residues from the Gɑ and Gβ subunits further stabilize the anchor 1 region (**Fig. 2c**), as shown by its very low B-factors (21.8 Å^2^), indicating an exceptional rigidity of the Gɑ and Gβ interface that assists to the stabilization of the heterotrimer.

We also observe a hydrophobic contact between the phenyl ring of FR and Ser53^G.H1.02^ in the P-loop of the GTPase domain (**Fig. 2d**). Ser53^G.H1.02^ is completely conserved in all G protein families and is involved in nucleotide binding and exchange by binding the phosphate groups of GDP/GTP and a Mg^2+^ ion. It has been shown that Mg^2+^ has a rather low affinity (in the millimolar range) to the GDP-bound state of G proteins^28, 29^ that increases significantly in the GTP-bound state towards a low nanomolar affinity. So, despite the presence of Mg^2+^ ions in the purification buffer at a concentration of 1 mM, we obtained a magnesium-free structure, as expected in the GDP-bound state. In our structure, the oxygen of the Ser53^G.H1.02^ side chain is oriented towards both phosphate groups of GDP. This places Ser53^G.H1.02^ between GDP and FR, resulting in a direct contact between the side chain and the inhibitor (**Fig. 2d**). Remarkably, this contact between the inhibitor and the P-loop was predicted by molecular dynamics simulations of Gq/FR complexes in which the relative mobility of the S53^G.H1.02^ side chain clearly differed between the apo and inhibitor-bound ensembles^30^. Here, we provide the first experimental evidence of a direct influence of FR on the nucleotide-binding pocket. We suspect that FR binding reduces the flexibility of the Ser53^G.H1.02^ side chain, which further hinders efficient nucleotide exchange, adding another feature to the inhibition mechanism of the Gq/11 family by cyclic depsipeptides.

Additionally, we observe another previously unidentified contact between FR and Tyr67^G.h1ha.04^ (**Fig. 2e**). This residue is part of linker 1 connecting the GTPase and α-helical domains, which is associated with the regulation of nucleotide exchange rates as well as GPCR interaction specificity^31, 32, 33^. So far, it was assumed that linker 1 showed reduced mobility in the presence of an inhibitor solely due to indirect interactions mediated by Arg60^G.H1.09 7^. The additional direct polar interaction with Tyr67^G.h1ha.04^ further stabilizes this crucial region. In addition, the structure shows that Arg60^G.H1.09 7^ and Tyr67^G.h1ha.04^ mutually stabilize each other through a cation-π-interaction (distance between the guanidinium moiety of Arg60^G.H1.09 7^ and the aromatic ring of Tyr67^G.h1ha.04^ = 4.5 Å).

Based on this high-resolution structure of the G11/FR complex, we have been able to identify several interactions not observed in the published Gq/YM structure^6^. However, it was not possible to determine whether these interactions (mediated by Arg96/Gβ, Ser53^G.H1.02^, and Tyr67^G.h1ha.04^) were unique to the binding mode of FR or if they could also be made by YM, as the lower resolution of the Gq/YM structure prevented precise modeling of the ligand binding pose. For instance, in the published Gq/YM crystal structure^7^, the inhibitor was not modeled close enough to Tyr67^G.h1ha.04^ for this interaction to be detected. Interestingly, there is unexplained positive *mF*_o_ − *DF*_c_ electron density in the Gq/YM maps exactly at this position, suggesting a conserved binding pose between FR and YM. We therefore decided to obtain a high-resolution structure of the G11/YM complex that would allow an unambiguous comparison of the binding modes of the two inhibitors.

We adapted the purification and co-crystallization protocols using YM instead of FR to grow crystals of a G11/YM complex under almost identical conditions. We solved the structure of the G11iN_3_-S:YM complex at a 1.7 Å resolution, allowing us to compare the binding modes of both ligands with a high level of confidence, revealing that the inhibitors FR and YM share a very similar binding mode (**Fig. 3**).

**Figure 3:**
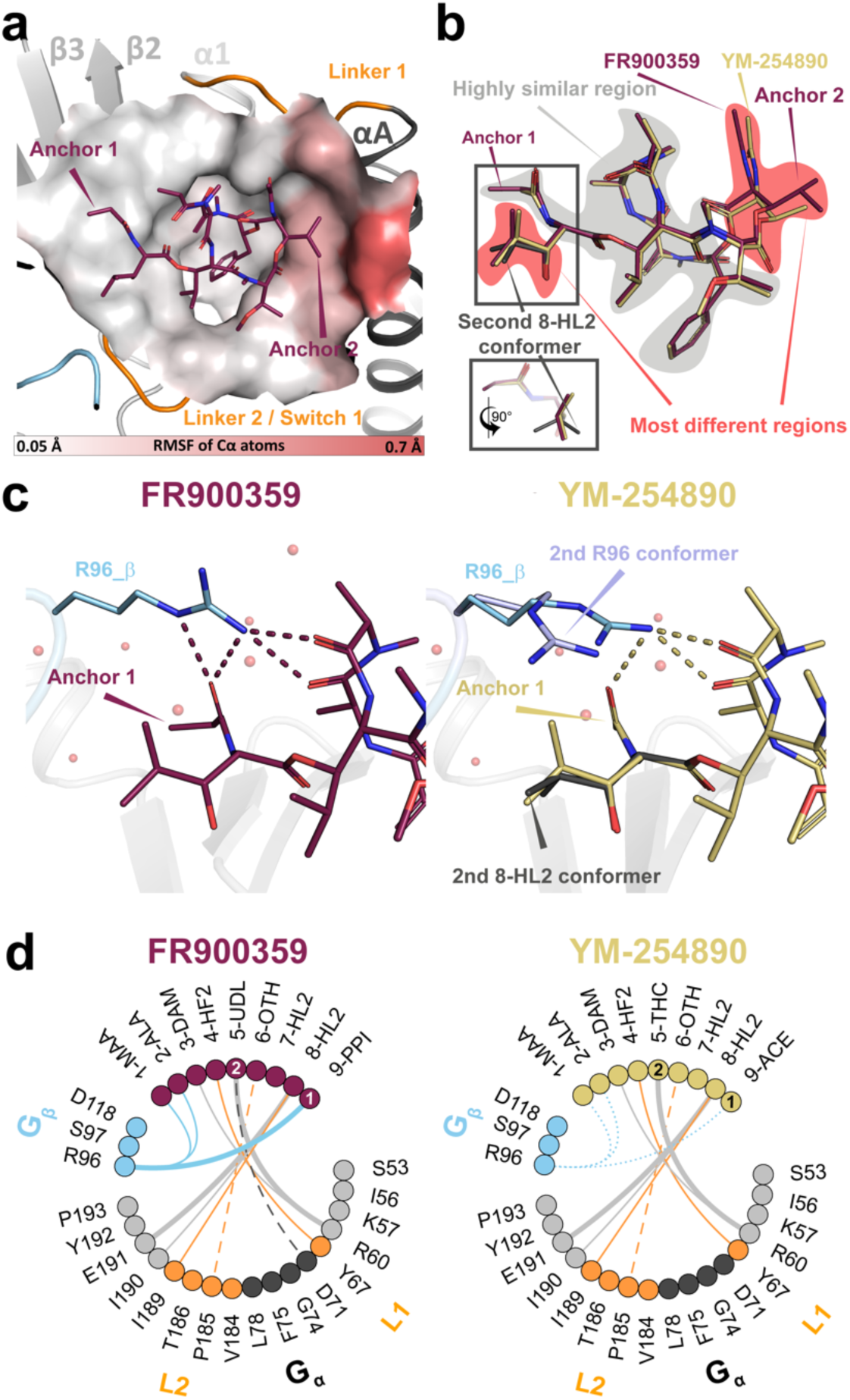
Comparison between the G11/FR and the G11/YM crystal structures. The structures were aligned on the Cα atoms of the protein. **a)** Local structural differences between the binding pockets (measured as relative root mean square fluctuation (RMSF) of the protein Cα atoms) are used to color the protein surface around FR. RMSF values are depicted with a color gradient from light grey (RMSF = 0.05 Å; nearly identical local protein structure between FR and YM) to red (RMSF = 0.7 Å; maximum differences). The changes are generally very subtle and can only be clearly observed in the region near anchor 2, where Asp71^H.HA.03^ and Gly74^H.HA.06^ of helix αA are mildly pushed away from FR compared to the G11/YM structure. **b)** Comparison of the ligand binding poses. The two ligands show only small structural differences. In particular, the ligand core (shaded in grey) is highly similar. The main differences appear near both anchors (shaded in red). A small difference in the position of anchor 2 in FR translates into a small local rearrangement of its backbone. In addition, a second conformer of the β-hydroxyleucine (residue ID: 8-HL2) side chain near anchor 1 is observed in YM with 0.5 occupancy. In the figure, residues in FR and YM are labeled according to their assigned residue ID **(Suppl.** Fig. 3). **c)** Detailed view of the region around anchor 1 of FR (left panel) and YM (right panel). The interactions of R96/Gβ with FR include four direct hydrogen bonds (red dashes). In the presence of YM, however, R96/Gβ displays two conformations: the first conformer (occupancy 0.5) is roughly similar to that of FR, forming three hydrogen bonds with YM (yellow dashes). Interestingly, a second conformer that is not contacting YM anymore is also observed. **d)** Interaction plots between both ligands and the binding pocket (FR left, YM right). Residues of FR/YM and of G11iN3-S within 4.5 Å of the ligands are represented as filled circles colored as in Fig. 2a. The lines depict contacts between residues within a radius of 3.5 Å. The thickness of the line represents the number of inter-atomic interactions between the residues (thin = single interaction; thick = two interactions). Solid lines correspond to polar interactions and dashed lines to van-der-Waals contacts. In the YM plot, the conformational variability of R96 is depicted by representing its weaker interactions with the ligand as dotted lines.

The two protein-ligand complex structures align perfectly well (RMSD = 0.207 Å), including the ligand-binding sites of FR and YM, which appear to be virtually identical. All newly described ligand-protein interactions of the G11/FR structure are also present in the G11/YM structure, proving a highly conserved mode of action between both inhibitors. However, we could identify small but crucial differences between the binding modes. To understand the structural determinants of FR and YM binding, we analyzed in detail the differences in the protein backbone position (**Fig. 3a**), the ligand-binding poses (**Fig. 3b**), as well as the protein side chain positions (**Fig. 3c**).

Only subtle changes in the protein backbone can be observed around anchor 2 (**Fig. 3a**), where Asp71^H.HA.03^ and Gly74^H.HA.06^ of helix αA are mildly pushed away from FR compared to YM. Both ligands also adopt a virtually identical pose in the binding pocket (**Fig. 3b**). Again, the main difference occurs at anchor 2, as the isopropyl moiety of FR is slightly shifted compared to the methyl group of YM due to a steric clash with Gly74^H.HA.06^. This displacement has only mild consequences on the structure of the cyclic backbone of FR, e.g., the N-acetyl side chain is relocated by 0.7 Å towards the opposite side of the pocket (i.e., towards anchor 1). Another change in the ligand-binding pose is the presence of a second conformer of the β-hydroxyleucine side chain near anchor 1 in YM where one methyl group is rotated by ∼80° (**Fig. 3b**).

The most noticeable difference between the G11/FR and G11/YM complexes is at Arg96/Gβ, the only contact between the inhibitors and the Gβ subunit (**Fig. 3c and 3d**). We observed an additional contact between FR and Arg96/Gβ, which is not present in the YM structure. Moreover, our data show a second conformer of Arg96/Gβ in the YM structure (**Fig. 3c**) that relocates the side chain away from YM to interact with Asp118 also in the surface of Gβ through a water molecule. Both Arg96 and Asp118 are highly conserved in Gβ subfamilies. One conformer (occupancy = 0.5) still interacts with YM but is tilted by ∼70° compared to the G11:FR structure resulting in the loss of one of the four hydrogen bonds to the inhibitor. Notably, the second conformer is located above the β-hydroxyleucine residue of YM **(8-HL2 in** **Fig. 3c**), indicating a possible coupling between these two elements. Thus, we hypothesize that the presence of FR results in increased stability of the Arg96 side chain and the Gα-Gβ interface compared to YM, suggesting an important role for this residue in the stabilization of the heterotrimer. At the other side of the binding pocket, anchor 2 of FR forms one more van-der-Waals contact to Asp71^H.HA.03^ **(Suppl. Fig. 4)** compared to YM.

In addition to direct interactions between ligand and protein, water-mediated effects can contribute significantly to binding^34^. The high resolution of our electron density maps allowed us to model a total of 760 and 804 water molecules in the FR/G11 and YM/G11 complexes, respectively. This allowed us to analyze the differences in the topology of the water structure in both ligand-binding sites in detail. Typically, enthalpically favorable (“happy”) water molecules can mediate protein-ligand interactions that represent a substantial contribution to the free binding energy. However, we do not observe any FR/YM interactions with the G protein-mediated by “happy” buried waters. In moderately hydrophobic ligands such as FR and YM (logP values of 1.86 and 1.37, respectively^15^), the displacement of poorly ordered (“unhappy”) water molecules from the also hydrophobic binding pocket^18^ can contribute appreciably to the ligand free binding energy.

In addition to these well-known effects, the formation of enthalpically favorable first-shell hydration layers around solvent-exposed parts of ligands has been identified to be relevant for binding^35^. Many of the water molecules in this layer do not mediate direct interactions between protein and ligand but form instead a stable network of water molecules surrounding the exposed parts of the ligand. The role of this layer is particularly consequential in shallow and open binding pockets exposed to the solvent, such as in the case of FR and YM (the calculated solvent-exposed surfaces of the ligands are 134 Å^2^ for FR and 129 Å^2^ for YM). We observed an extended and continuous hydrogen bonding network of first-shell waters that extends to polar functional groups of the ligand and the protein (**Fig. 4** **and Suppl. Table 2**). Even though many of these waters do not directly contact the ligand, they all show strong electron densities, indicating that they are well ordered (**Suppl. Fig. 5a**). Particularly, the water chain H-I-J spanning over YM shows strong signals, indicating high stability. Remarkably, this central element of the water network is absent in the FR complex. Overall, the electron densities in the first-shell water layer are weaker in FR than in YM, indicating a relative desolvation of FR compared to YM (**Fig. 4**).

**Figure 4:**
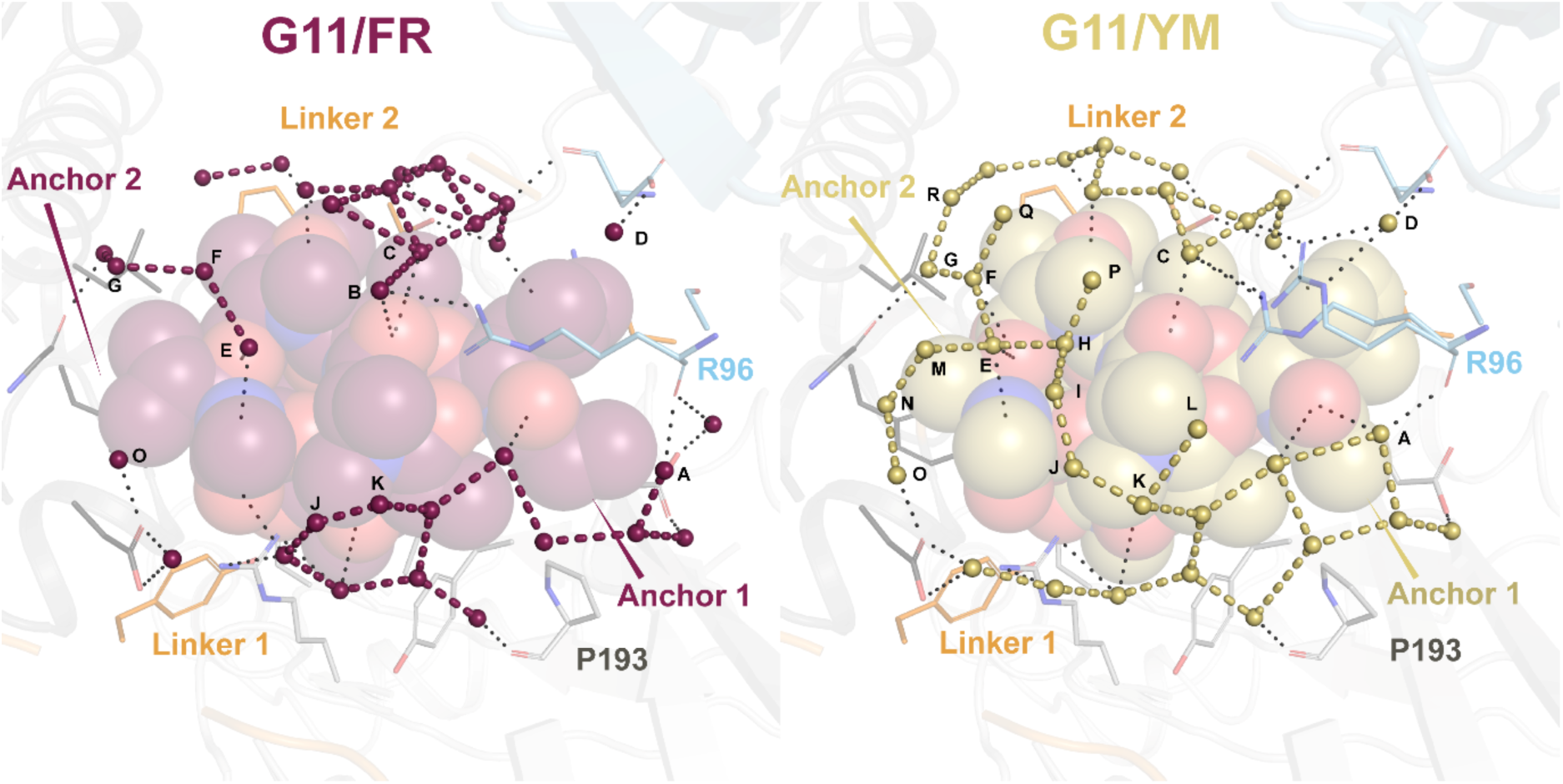
Ligand solvation. Water networks in the first solvation shell of FR (left panel) and YM (right panel). Water molecules are represented as small spheres (FR = red, YM = dark yellow), ligands are shown as spheres (FR = red, YM = dark yellow), the interacting protein residues as sticks, and the protein backbone as translucent cartoons. Water-water interactions are indicated with thick dashes and water-ligand and water-protein interactions are represented by thin black dashes. Differences in the solvation of both ligands, with a stronger solvation of YM, could potentially contribute to the different binding profiles of FR and YM.

There are some differences in the local structure of the first-shell waters near anchor 2 of the inhibitors. In YM (**Fig. 4** **right panel**), waters M and N connect to the central water E. In FR, the steric hindrance and increased hydrophobicity of the isopropyl moiety of anchor 2 displaces these two waters. Other waters in this region are apparently displaced as a result of subtle but significant differences in the binding poses of FR and YM around anchor 2 (**Fig. 3b**). In the more structured first-shell layer of YM, water E is coordinated via two hydrogen bonds to YM and acts as one of the central nodes of a dense water network formed by waters E, P, H, I, and J (**Fig. 4** **right panel**). In FR, anchor 2 is moderately pushed away from helix αA of the Gα subunit, resulting in a small change (0.7 Å) at the FR backbone at the N-acetyl side chain of the N-acetyl-hydroxyleucine (**Fig. 3b** **and Suppl. Fig. 5c**). Water E follows this relocation of the backbone carbonyl destabilizing the central water network P-H-I-J while still maintaining the coordination with the ligand. Interestingly, anchor 2 has been proposed to be crucial for the high affinity of FR^20^.

At the other side of the inhibitors, near anchor 1, the first-shell layer over the ligands is less dense due to the shielding by Arg96 of Gβ. The smaller acetyl group in anchor 1 of YM allows the presence of water A that bridges the inhibitor to the backbone of Arg96 (**Fig. 4** **right panel and Suppl. Fig. 5b**). However, the larger propionyl in anchor 1 of FR does not allow the presence of water A (**Suppl. Fig. 5b**), resulting in the relocation of the Arg96 side chain closer to anchor 1 of FR. This allows the formation of further polar and hydrophobic interactions with FR, which may translate into the observed conformational stability of the Arg96 side chain.

In summary, our structures reveal three possible mechanisms for heterotrimer stabilization by FR and YM. The inhibitors (i) contact the P-loop of the GTPase domain, (ii) reduce the rigidity of linker 1, and (iii) the long side chain of Arg96 in Gβ disrupts a first-shell layer of waters in a shallow binding pocket and “clamps” over the inhibitors, tightly over FR and more loosely over YM. We hypothesized that this direct interaction with the Gβ subunit could translate into an enhanced physical interaction of Gα with Gβγ and hence stabilization of the G protein heterotrimer.

To test this hypothesis, we used site-directed mutagenesis to mutate Arg96 in Gβ to alanine and then compared the thermal stability of wild type and R96A heterotrimer in the presence and absence of the inhibitors (**Fig. 5**). In the apo state (no inhibitors), the wild type and mutant constructs display near-identical biphasic curves (**Fig. 5a**). Addition of FR or YM results in a significant thermal stabilization and a monophasic unfolding behavior. In the wild type, the curves for both ligands show an identical onset of the unfolding event; however, the curves decouple, and YM results in a steeper denaturation curve. Interestingly, this feature disappears almost entirely in the R96A mutants. For clarity, the melting temperatures are shown in **Figure 5b** as bar graphs. In the absence of inhibitors, the mutation has no significant effect on stability. In contrast, upon ligand-complex formation, mutation of Arg96 results in a noticeable destabilization for both ligands (**Fig. 5c**), highlighting the functional role of the interaction of the two inhibitors with the Gβ subunit.

**Figure 5:**
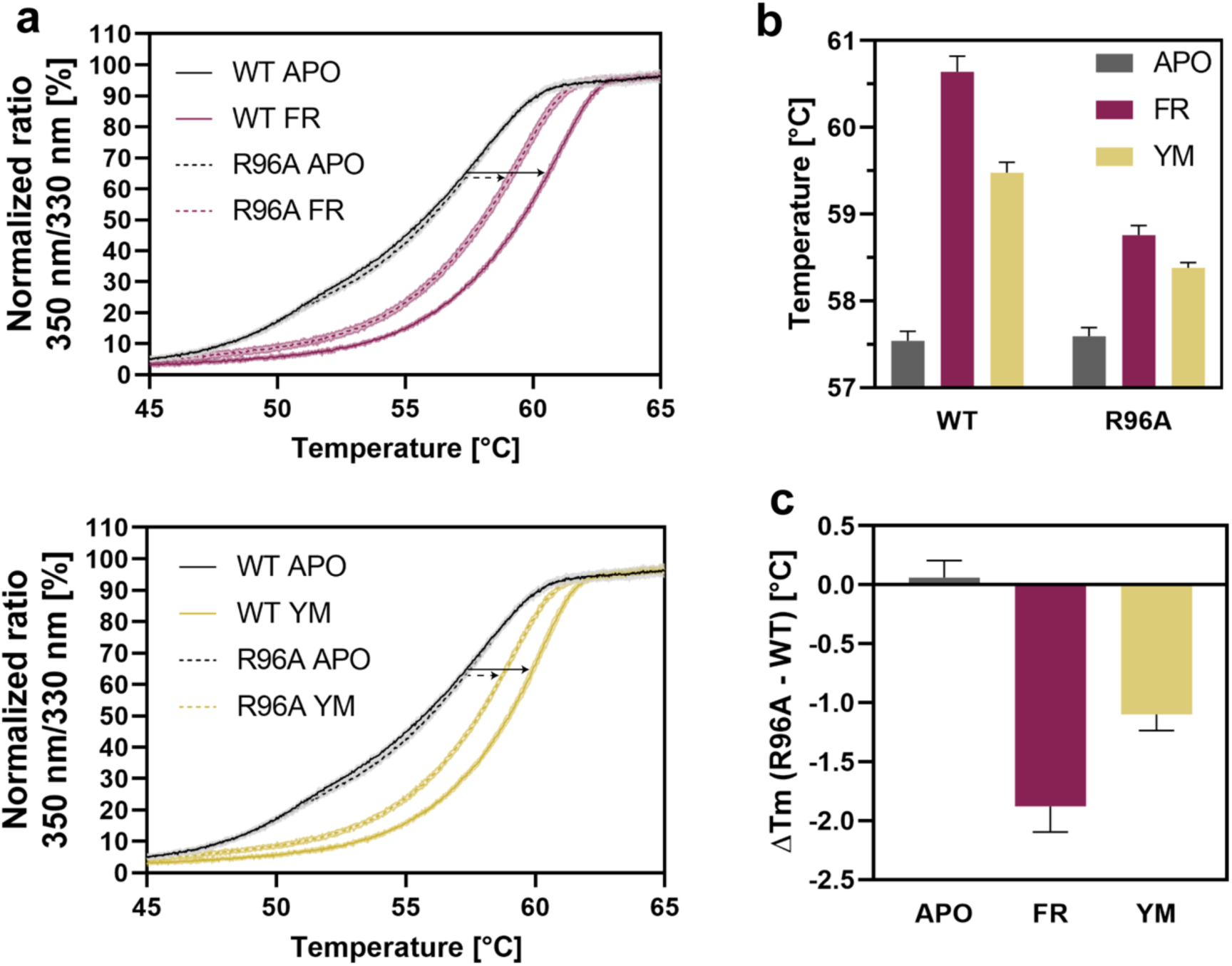
Destabilizing effect of the R96A mutation. **a)** Protein unfolding curves determined by the red shift of tryptophan fluorescence (compare to Fig. 1b). The curves are shown as solid lines for G11iN3-S WT and as dashed lines for G11iN3-S with the R96A mutation in the Gβ subunit. The lines are colored black (absence of inhibitor), red (in the presence of FR, top), and dark yellow (in the presence of YM, bottom). The arrows indicate the temperature shift upon ligand addition. **b)** Calculated melting temperatures (Tm) represented as a bar graph. Apo samples are shown in grey (using the second unfolding event as a reference), FR in red, and YM in dark yellow. **c)** The calculated ΔTm (Tm (R96A) – Tm (WT)) highlights the overall destabilization induced by the mutation and the stronger effect in the FR-bound complex: ΔTm (FR) = -1.88 ± 0.22 °C; ΔTm (YM) = -1.10 ± 0.10 °C.

To investigate the functional relevance of the interaction between inhibitor and Gβ in a cellular environment, we monitored the effects of FR and YM on Gα11β1γ2 heterotrimers in their basal state in living cells. We restrained the inhibitor effects to Gα11 heterotrimers by re-expressing Gα11 (wildtype or mutant) in CRISPR/Cas9 gene-edited HEK293 cells lacking all other Gα subtypes apart from Gαi/o (HEK293 Δ7)^36^. We then employed a bioluminescence resonance energy transfer (BRET)-based approach measuring the competition for Gβγ binding between the Gα11 subunit and a membrane-associated Gβγ-effector mimic, the C-terminus of GPCR kinase 3 (masGRK3ct). BRET was conceived by tagging the Gβγ dimer with a split Venus as BRET acceptor and masGRK3ct with nano luciferase as the BRET donor. In this experimental setup, sequestering of free Gβγ dimers by masGRK3ct produces high BRET signals, which are diminished by co-expressed Gα11. A stabilization of Gα11Gβγ heterotrimers by FR and YM would shift the equilibrium towards Gα-bound Gβγ and away from Gβγ:masGRK3ct, and hence manifest in a BRET decrease (assay principle, **Fig. 6a**).

**Figure 6:**
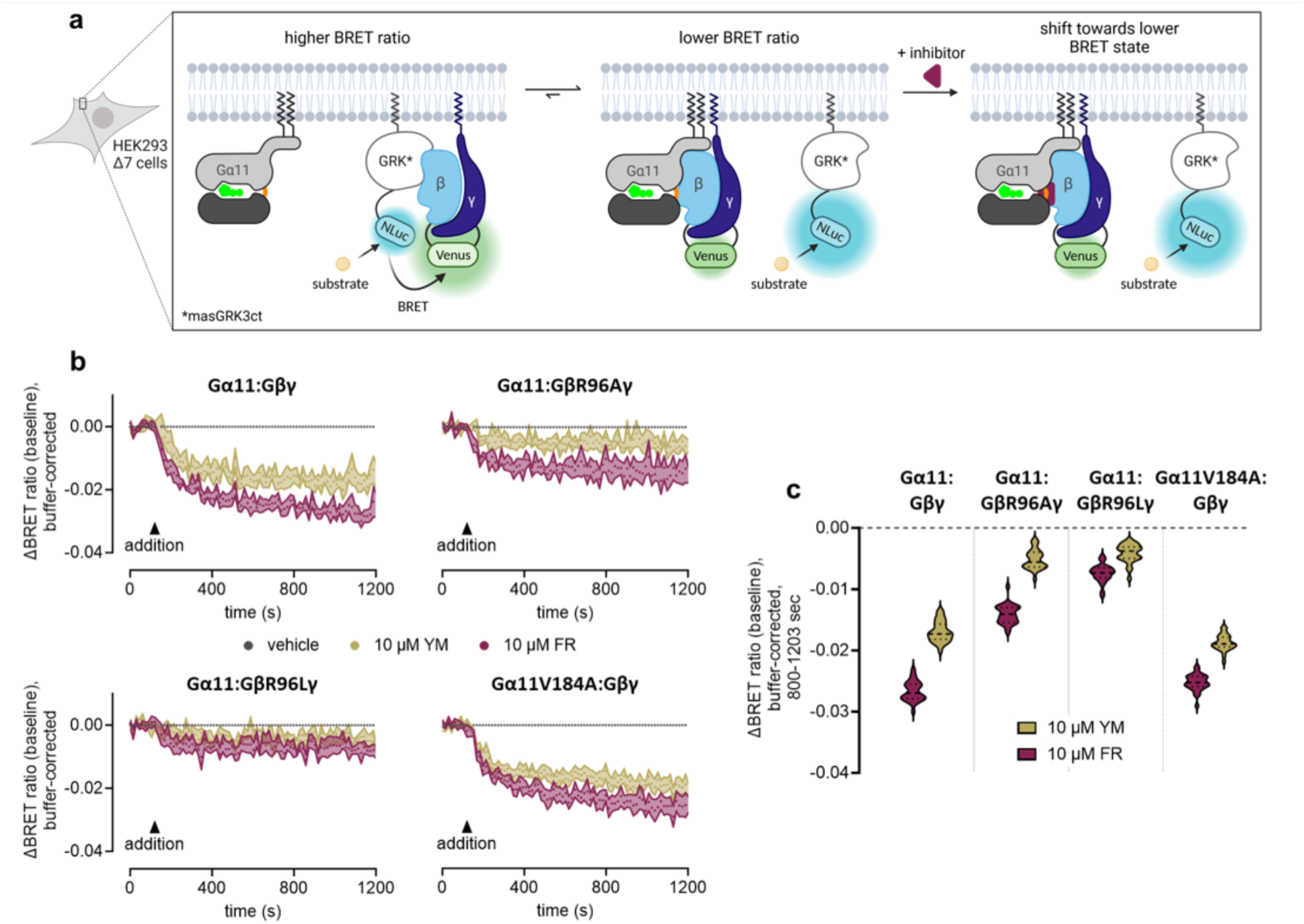
FR and YM stabilize the Gα11:Gβγ heterotrimer through an interaction with R96/Gβ. **a)** BRET assay in living cells, in which Gβγ is tagged with split Venus (BRET acceptor) and masGRK3ct with nano luciferase (Nluc, BRET donor). The binding of free Gβγ to masGRK3ct results in a basal BRET signal. Inhibitor-induced heterotrimer stabilization shifts the equilibrium further towards Gα-bound Gβγ resulting in a decrease in the signal. **b)** Effect of FR or YM addition (10 µM, saturating concentration) on the BRET signal in HEK293 Δ7 cells transiently transfected with the indicated Gα11 and BRET sensor constructs, as well as PTX-S1. **c)** Violin plot summary of the BRET data. In wild type G11, both FR (red) and YM (dark yellow) induced a decrease in the BRET ratio indicating a stabilization of the G protein heterotrimer. Mutation of R96/Gβ to alanine or leucine reduced this effect for both inhibitors. A mutation in Gα11 at the other side of the ligand binding pocket (V184A) does not alter the inhibitor-induced effect. BRET traces are mean + SE of at least five independent biological replicates.

The addition of both FR and YM detectably decreased the basal BRET ratio, indicating a shift of the G protein population towards Gαβγ heterotrimers (**Fig 6b; top left panel and** **Fig. 6c**). However, though likely, it is not clear whether the observed BRET decrease invariably depends on a direct physical interaction between FR an YM and arginine at position 96 in Gβ. To investigate a causal link for the role of Arg96/Gβ in inhibitor-mediated heterotrimer stabilization, we replaced this residue with alanine – to remove all Gβ interactions to the inhibitor– and with leucine –to solely eliminate the polar Gβ interactions to the inhibitor but maintain those mediated by the hydrophobic part of the Arg side chain in the wild type Gβ (**Fig. 2c**). This chosen experimental setup is particularly suited to interrogate a direct interaction between FR and YM and the Gβγ subunit complexes because tagging of Gβγ by split Venus allows to invariably assign BRET changes to the nature of the transfected Gβ subunit, thereby bypassing the confounding contribution of endogenous Gβ proteins.

Both mutations of Arg96 significantly impacted the stabilizing effect of the inhibitors on the heterotrimer in unstimulated cells, with a more prominent effect in the leucine mutant (**Fig. 6; top right and bottom left panels and** **Fig. 6c**). To support the specific role of the Gβ subunit in heterotrimer stabilization and exclude possible long-range allosteric effects involving Gα, we introduced a mutation at the opposite side of the inhibitor binding site, V184^G.hfs2.03^ in linker 2/switch I. While this residue is known to contribute to the inhibitory action of FR and YM on Gαq^17^, its mutation to alanine does not dampen heterotrimer stabilization by the inhibitors (**Fig. 6b; bottom right panel and** **Fig. 6c**), as was the case for the Gβ mutants. Neither of the mutations impacted activation of G11 heterotrimers by carbachol-stimulated muscarinic acetylcholine receptor 3 after exogenous transfection or the inhibitory potency of FR and YM, attesting intact activation and inhibition profiles comparable to those observed with G11 wild type proteins **(Suppl. Fig. 6)**.

In summary, the BRET data supports the stabilization of the entire heterotrimer –i.e., not only of the Gα subunit– as a molecular feature that may contribute to the observed efficient suppression of Gq/11 signaling by FR and YM. Unexpectedly, Gβ rather than Gα appears to play a central role in mediating this stabilizing effect. However, because mutation of R96/Gβ does not impact inhibitor potency, our data also indicate that inhibitor:Gα interaction is sufficient to block Gα activity.

Taken together, our results provide an in-depth insight into how FR and YM, two highly specific inhibitors, shut off Gq/11 signaling in mammalian cells. Stabilization of the flexible hinge regions and constraint of Ser53^G.H1.02^ in a position that is not compatible with activation stabilize Gα in an inactive GDP-bound conformation. The direct contact of the inhibitors to Gβ enhances the stability of the full heterotrimer, securely locking the G protein in its unproductive heterotrimeric form and preventing the conformational changes leading to nucleotide exchange and dissociation necessary for activation.

## DISCUSSION

Our structures expand our understanding of how the FR and YM depsipeptides inhibit Gq/11 proteins. We propose that the inhibitors securely lock Gq/11 protein heterotrimers –and not only the Gα subunit– by engaging Gβ. We provide three lines of experimental evidence independently supporting the role of FR and YM in G11 protein heterotrimer stabilization. First, our high-resolution crystal structures reveal that the inhibitors join Gα and Gβγ together, with Arg96 in Gβ acting as a connector that clamps over both inhibitors (**Fig. 3c and d**). Second, *in vitro* thermostability assays reveal that the inhibitors enhance the stability of Gα:Gβγ complexes (**Fig. 1b and 5**). Third, cellular BRET assays show this effect in living mammalian cells. Using site-directed mutagenesis, we could confirm that the contact between Gα and Gβγ mediated by the interaction of the inhibitor with Arg96 decisively contributes to the stabilization of the G protein heterotrimer (**Fig. 5 and 6**). Remarkably, a mutation at Arg96 in human Gβ1 has been implicated in affecting the assembly of G protein heterotrimers leading to global developmental delay^37^. We propose that the FR and YM depsipeptides can be thought of as ‘adhesives’ that bind together not only the GTPase and α-helical domains of Gα but the entire heterotrimer including Gβ. This leads to the formation of a completely ‘silent’ complex unable to engage in cellular signaling pathways.

Our crystallographic data shows that the structures of Gα11 and Gαq are virtually identical, as expected due to their high degree of sequence identity **(Suppl. Fig. 7)**. Also, as foreseen in previous studies, we show that FR and YM bind in the same pose, with only subtle rearrangements in the protein backbone at the N-terminal end of helix αA in Gα (near anchor 2 of the inhibitors) and in the side chain of Arg96 in Gβ (near anchor 1) (**Fig. 2**). The lack of a reference structure of a “free” Gαq/11 heterotrimer (i.e., not bound to an inhibitor or an effector) impedes full appreciation of the inhibition mechanism, as the impact of ligand binding on the overall protein conformation remains elusive.

While the binding modes of FR and YM are extremely similar, they are not identical. YM, with smaller anchors 1 and 2, seems to fit better in the shallow binding pocket, in which Arg96 in Gβ exists in two equally populated conformers establishing only transient interactions with the inhibitor. A persistent layer of first-shell water molecules over the solvent-exposed part of YM, in addition to the probable displacement of poorly ordered waters in the hydrophobic shallow binding pocket, likely contribute to its good binding affinity (pKi = 8.23)^20^. On the other hand, the larger ethyl in anchor 1 of FR alters the first-shell water layer, facilitating the formation of strong direct hydrogen bonds to Arg96. This enthalpy gain may explain the higher binding affinity (pKi = 9.23)^20^ and the stronger stabilization of the heterotrimer by FR (**Fig. 5 and 6**).

The remarkable order of the first-shell waters can be attributed to the rigidity of the ligands in the binding site, as evidenced by their well-defined electron density **(Suppl. Fig. 2)**. NMR studies have shown that the cyclic depsipeptide backbones of FR^22^ and YM^38^ also exist primarily in a well-defined conformation in water, with all amide bonds in a *trans* conformation except for one *cis* amide bond between the phenyllactic acid and the N-methyl-α-β-dehydroalanine residues, which are the same conformations that we observe in our structures. This rigidity of the depsipeptide backbone is compatible with a conformational selection binding mechanism^26^ in which the reduced accessible conformational space constrains the ligand close to a bioactive conformation that only requires small rearrangements in the protein (e.g., in the N-terminal end of helix αA in Gα for FR, **Fig. 3a**) for productive binding. There may be greater diversity in the ligand side chain conformations, as observed in the β-hydroxyleucine (residue ID: 8-HL2) side chain near anchor 1 of YM (**Fig. 3b**). In this case, the preferred conformation will depend on the precise backbone geometry but also on the interaction of the ligand side chains with neighboring residues in the peptide or the protein. Thus, there may still be a part of induced fit in the ligand binding mechanism.

Our structures allow the rationalization of observed structure-activity relationships of FR and YM synthetic derivatives^20, 22^. Essentially, most modifications drastically reduce their inhibitory ability, whereas the majority of natural analogs, which mostly present variations at the acyl chain of anchor 1, are still active, albeit with a reduced affinity to Gq and inhibitory activity (e.g., the non-natural analog FR5, which harbors a larger butyryl moiety in anchor 1)^9^. Thus, there seems to be some room for structural variation at the acyl chain (one of the critical determinants for optimal interaction of the inhibitors with Arg96/Gβ and, consequently, for efficient inhibition of the heterotrimer) but not in most other parts of the molecule. Interestingly, anchor 1 of FR is directed towards a polar pocket (Ser98 in Gβ and Arg202^G.S3.04^ in Gα) that includes several well-ordered water molecules **(Suppl. Fig. 5b)**, hinting at possible ways to derivatize these inhibitors (e.g., by adding a polar group at anchor 1) to further stabilize the Gα-Gβ-ligand interface. The FR isopropyl group of anchor 2 (contacting Asp71^H.HA.03^ of the Gα subunit) could also be derivatized to establish additional hydrogen bonds to the protein, e.g., by replacing it with a polar moiety. We anticipate that our high-resolution structures will contribute to the targeted development of new FR derivatives by structure-guided drug design and fragment screening.

Interestingly, there is another example of heterotrimer stabilization as an efficient mechanism to switch off G proteins. The antibody fragment FAB16^39^ slows down nucleotide recruitment to Gαi and prevents heterotrimer dissociation upon treatment with GTPγS by recognizing an interface between the Gα and Gβγ subunits. However, unlike FR and YM, FAB16 is not selective for the nucleotide-binding state of the G protein. This suggests that the entire Gα-Gβ interface of G protein heterotrimers could be the target of structure-based drug screening to find new inhibitors acting as “Gα-Gβ clamps” that hinder heterotrimer dissociation. We foresee that our results will advance our understanding of the G protein inhibition mechanism of macrocyclic depsipeptides and how they effectively blunt mammalian Gq/11 signaling in living cells.

## Data availability

Coordinates and structure factors have been deposited in the Protein Data Bank under accession numbers 8QEG and 8QEH. Source data and supplementary data files are provided in this paper. All other relevant data supporting the key findings of this study are available within the article, its supplementary information, or from the corresponding authors upon reasonable request.

## Acknowledgements/Funding

The authors thank the Deutsche Forschungsgemeinschaft (DFG, German Research Foundation) – Project number 290827466/FOR2372 (grants CR464/7-1 to M.C., KO 902/17-2 to G.M.K., 290847012/FOR2372 to E.K., 216619161/FOR2372 to G.S., and 418513893/FOR2372 to X.D.) and the Swiss National Science Foundation – Project numbers 192780 (to X.D.) and 183563 (to G.S., under the Sinergia program) for funding. M.J.R. received funding from the European Union’s Horizon 2020 Research and Innovation Programme under the Marie Skłodowska-Curie grant agreement No. 701647. We thank the SLS beamline scientists for their support during crystallography data collection.

## Author contributions

X.D., G.S., and E.K. conceived, designed, and coordinated the research. J.M. performed protein expression, purification and functional characterization, crystallization, and crystal screening. M.J.R. and J.M. collected and processed the X-ray data. J.A., L.J., and H.S. designed, performed, and analyzed the cellular BRET assays. A.B. and F.A. performed and analyzed the grating-coupled interferometry assays. M.C. and G.M.K. provided FR. J.M., M.J.R., J.A., L.J., R.G-G., H.S., E.K., G.S., and X.D. interpreted the data. J.M. and X.D. wrote the manuscript with the assistance of M.J.R., J.A., M.C., G.M.K, M.H., E.K., and G.S.

## MATERIALS AND METHODS

### Chemicals

FR900359 was isolated and purified from *Chromobacterium vaccinii* as described elsewhere^9, 40^ YM-254890 was purchased from Fujifilm.

### Construct design for crystallography

**Gα11iN1-29**: a Gα11 chimera with the N-terminal helix of Gαi_1_ was constructed as described elsewhere^25^. In addition, an N-terminal StrepII-eGFP-tag followed by a 3C protease cleavage site was added to improve expression and purification strategies. The full gene was ordered from GeneWiz.

**Gβ1**: human Gβ1 was N-terminally tagged with a deca-histidine tag followed by a 3C protease cleavage site. The R96A mutant was ordered from GeneWiz.

**Gγ2 C68S**: Residue C68 was replaced with a serine by site-directed mutagenesis of human Gγ2 to remove the prenylation site and make the G protein trimer soluble.

All three subunits were cloned into one pAC8REDNK backbone^41^ (kind gift of Dr. Arnaud Poterszman, Strasbourg) under the regulation of individual PH or P10 promoters.

**GST-Ric8A**: The expression chaperone Ric8A was N-terminally fused to GST followed by a TEV protease cleavage site as described elsewhere^42^. The full-length gene was synthesized and cloned into a pAC8REDNK backbone by GeneWiz.

### Virus production and protein expression

Baculo viruses were produced in Sf9 insect cells using the *FlashBac* technology (Oxford Expression Technologies, Oxford, UK). 1.0 x 10^6^ Sf9 cells were co-transfected with 1.5 μg plasmid DNA and 500 ng linearized Bac10:KO1629 viral DNA using Cellfectin II (Invitrogen) following the manufacturer’s protocol. Cells were incubated for 5 hours at 27 °C with the transfection mixture before it was replaced by fresh SF900II SFM insect cell medium (Invitrogen) supplemented with 1% penicillin/streptomycin (PAN Biotech, Aidenbach, Germany). After 6-7 days of incubation at 27 °C, V_0_ generation virus was harvested by centrifugation at 800 x g and 20 °C, the supernatant supplemented with 1% Pen/Strep and 10% FCS (Sigma-Aldrich, Missouri, USA) and stored in the dark at 4 °C.

Viruses were two times amplified by infecting new Sf9 cultures at 2.0 x 10^6^ cells/ml in SF900II SFM medium with 1% V_0_ or V_1_ generation viruses, respectively. V_1_ or high-titer V_2_ generation viruses were harvested 72 hours post-infection by centrifugation as described above. However, only 1% FCS was used as a supplement.

The G11 heterotrimer was expressed in High Five insect cells (Invitrogen, Waltham, USA) by co-infection with a virus encoding for all three subunits and a virus encoding for GST-Ric8A. Optimal relative virus concentrations were determined in small-scale analysis by Fluorescence-detection Size Exclusion Chromatography^43^. Large-scale expression was conducted in High Five insect cells at a density of 4.0 x 10^6^ cells/ml in SF900II SFM medium using 5L Erlenmeyer flasks (Corning, Glendale, Arizona). Prior to expression, the medium was exchanged, and cells were resuspended at 4.4 x 10^6^ cells/ml in SF900II SFM medium. Then both viruses were added in an optimal ratio and the cells diluted to 4.0 x 10^6^ cells/ml. Cells were cultured at 27 °C and 120 rpm and harvested 48 hours post-infection by centrifugation. Pellets were flash-frozen in liquid nitrogen and stored at -80 °C.

### Protein purification

The G11iN_3_-S WT and R96A heterotrimers were purified on ice or at 4 °C as described elsewhere^7^ with small modifications. Frozen cell pellets were thawed in cold water and resuspended in Lysis Buffer (20 mM HEPES pH 7.5, 100 mM NaCl, 20 mM Imidazole pH 7.5, 3 mM MgCl_2_, 100 µM EDTA, 5 mM β-Mercapoethanol, 20 µM GDP, Roche cOmplete protease inhibitor cocktail and DNAse I) in a 1:1 (v:v) ratio. Cells were lysed in 3 cycles at 8000 PSI using an LM10 Microfludizer (Microfluidics International Corporation, Westwood, USA), and the lysate was clarified by ultracentrifugation at 186,000 x g for 45 min. The supernatant was incubated with TALON resin (Takara Biotech, Saint-Germain-en-Laye, France) for 90 min in a ratio of 1 ml resin per 10 g of cells. Beads were washed in batch mode with 3 x 10 CVs lysis buffer, followed by 2 x 10 CVs washes in TALON Buffer A (20 mM HEPES pH 7.5, 100 mM NaCl, 20 mM Imidazole pH 7.5, 1 mM MgCl_2_). The resin was resuspended in another 10 CVs of TALON Buffer A and transferred to a BioRad gravity flow column. A high salt gravity flow wash was conducted with 10 CVs TALON Buffer B (TALON Buffer A + 300 mM NaCl) before the G11iN_3_-S heterotrimer was eluted from the resin with 2 x 2 CVs TALON Buffer C (TALON Buffer A + 300 mM Imidazole pH 7.5) by gravity flow filtration. All tags were cleaved off overnight with homemade HRV 3C protease in a 1:100 (w:w) ratio while dialyzing against 70 CVs Dialysis Buffer (TALON Buffer A with 100 µM TCEP instead of β-mercaptoethanol). On the next day, precipitated salts were removed by centrifugation (10 min, 3200 x g), the supernatant supplemented with Imidazole pH 7.5 to a final concentration of 20 mM and incubated with TALON resin for 60 min. Flow-through was collected and the resin was washed with 2x 1 CV Dialysis buffer. All fractions were pooled and loaded onto a pre-equilibrated HiTrap Q HP 5 ml column (Cytiva, Marlborough, USA). The column was subsequently washed with 10 CVs HiTrap Q buffer A (same as Dialysis buffer) and the protein eluted with a linear gradient to 50% HiTrap Q buffer B (HiTrap Q buffer A + 1000 mM NaCl) over 20 CVs. The column was then washed with 100% HiTrap Q buffer B. Residual impurities of eGFP were removed on a HiLoad Superdex 200 pg 16/600 (Cytiva, Marlborough, USA) pre-equilibrated in SEC buffer (Dialysis buffer + 400 µM GDP). Fractions were analyzed on a 4-20% MiniProtean TGX SDS-PAGE (BioRad, Cressier, Switzerland). The gel was stained with Der Blaue Jonas (German Research Products, Haag a.d. Amper, Germany), and relevant fractions were pooled and concentrated to > 36 mg/ml (416 µM) using an Amicon Ultra centrifugal filter with a 50 kDa MWCO (Merck Group, Darmstadt, Germany).

### Protein crystallization

Concentrated G11iN_3_-S heterotrimer solution was supplemented with a 2-fold molar excess of FR900359 (100 mM stock in DMSO), incubated for 30 min, and diluted with SEC buffer to a final concentration of 32 mg/ml (370 µM) using an Amicon Ultra centrifugal filter with a 50 kDa MWCO (Merck Group, Darmstadt, Germany). The protein solution was used directly for protein crystallization or stored for up to 10 days on ice. Prior to crystallization, the protein solution was centrifuged for 30 min at 21k x g. Commercial sparse-matrix screens were set up in MRC-2 plates (SWISSCI, Neuheim, Switzerland) using a Mosquito (SPT Labtech, Melbourn, UK) with drops consisting of 200 nl protein solution + 200 nl reservoir solution at 4 °C and 20 °C. Several initial hits were obtained at both temperatures from the ProPlex screen (Molecular Dimensions, Rotherham, UK). The crystals were optimized in several rounds of alternating grid screening and additive screening using the Additive screen (Hampton Research, Aliso Viejo, USA) and MemAdvantage (Molecular Dimensions, Rotherham, UK) based on condition ProPlex H7. Final rod-shaped G11iN_3_-S-FR900359 crystals started growing after 14 days from heavy precipitate and continued to grow over 2 months at 4 °C in MRC-2 plates in a precipitant solution containing 0.09 M Na Acetate pH 4.5, 2.7% PEG Smears Medium, 6.3% MPD, 0.5 n-octyl-β—D-Glucoside and 10 mM Zinc sulfate heptahydrate. G11iN_3_-S-YM-254890 crystals grew within 3 days at 4 °C in MRC-2 plates in a precipitant solution containing 0.09 M Na Acetate pH 4.5, 2.7% PEG Smears Medium, 6.3% MPD, 0.5 n-octyl-β—D-Glucoside and 3% D-Trehalose.

### Data collection and processing

X-ray diffraction data were collected at the Swiss Light Source, Paul Scherrer Institute, Villigen, Switzerland, on beamlines PXI and PXIII. Crystals of G11iN_3_-S:FR900359 and G11iN_3_-S:YM-254890 belonged to the space group P 2_1_2_1_2_1_ and diffracted to 1.43 Å and 1.7 Å resolution, respectively. Datasets were processed with autoPROC^44^, which performs integration with XDS^45^ and scaling and merging with AIMLESS^46^. The G11iN_3_-S:FR900359 structure was solved by molecular replacement using PHASER^47^ with a publicly available Gq-YM complex structure^7^ (PDB code 3AH8) with ligand and water molecules removed used as a molecular replacement model. This model was later used to solve the molecular replacement solution for G11iN_3_-S-YM-254890. Both structures were interactively corrected and rebuilt in COOT^48^ and refined with REFMAC5^49^ in iterative cycles. Ligand restraints were generated with GRADE^50^. The quality of the structures was assessed with MOLPROBITY ^51, 52^ and PDB-REDO^53^. The data collection and refinement statistics are presented in Supplementary Table 1.

### GTPase Glo assay® (Promega, Madison, USA)

Reactions were set up in 384-well, low-volume, white, solid-bottom plates (Greiner Bio-One, Kremsmünster, Austria) and carried out at 20 °C under dim-red light conditions. Each condition was prepared with N = 8. 2.5 µL of protein (+/-FR900359) solution containing 2 µM G11iN_3_-S, 0.2 µM dark-state or light-activated Jumping Spider Rhodopsin 1^54^ were mixed with 2.5 µL of 2 µM GTP solution. The plate was sealed and incubated for 180 minutes shaking at 500 rpm. Subsequently, 5 µl of GTP-Glo reagent was added to each well and the reaction was carried out for 30 minutes. Finally, 10 µl of Detection Reagent was added. Luminescence was measured after 5 minutes in a PheraStar FSX (BMG labtech, Ortenberg, Germany) plate reader (Optical module: LUM plus, Gain: 3600, focal height:14, 1 second/well).

### Differential scanning fluorimetry

Melting temperatures were assessed using Differential Scanning Fluorimetry (DSF) on a Prometheus Panta (NanoTemper Technologies GmbH, Munich, Germany). Purified G11iN_3_-S WT and G11iN_3_-S R96A were diluted in DSF buffer (20 mM HEPES pH 7.5, 100 mM NaCl, 1 mM MgCl_2_, 50 µM GDP, 100 µM TCEP) to 5 µM in the presence of a 5-fold molar excess of FR or YM or an equivalent volume of DMSO. Mixtures were incubated for 15 min on ice. Samples were measured 33 times/ °C in a linear temperature gradient ranging from 35 °C – 80 °C with a slope of 0.5 °C/min. The optimal excitation energy was determined by auto-detection and set to 60%.

### Grating-coupled interferometry (GCI)

Kinetic binding measurements were performed using Grating-Coupled Interferometry (GCI) on a Creoptix WAVEdelta system (Creoptix-a Malvern Panalytical brand). Two flow channels of a PCH Creoptix WAVEchip (sensor chip) were conditioned according to the manufacturer’s specifications. The surface was activated by a 420-second injection of 0.4M EDC (1-ethyl-3-(3-dimethylaminopropyl)carbodiimide hydrochloride) and 0.1M NHS (N-hydroxysuccinimide) at a flow rate of 10 μl/min. Purified G11iN_3_-S was injected at 20μg/ml in immobilization buffer (10mM Na-Acetate, pH4.5) onto one flow channel and covalently coupled to a final level of 8000 pg/mm^2^. The surface of both channels was passivated by a 420-second injection of 1M Ethanolamine pH8 and then stabilized with several injections of running buffer (20 mM Hepes pH7.5, 100mM NaCl, 1mM MgCl_2_, 20 μM GDP, 100 μM TCEP, 2% (v/v) DMSO). FR was diluted from a 10 mM DMSO stock to 5 μM in running buffer and injected using the waveRAPID method^55^ with 6 pulsed injections of increasing duration over 120 seconds, followed by 5000 seconds of running buffer to follow FR dissociation from G11iN_3_-S. Data analysis was performed using the Creoptix WAVEcontrol software (Creoptix – a Malvern Panalytical brand). The response signals were double referenced, subtracting the signal recorded on the reference channel without protein as well as the signal recorded from blank injections with running buffer. Double-referenced signals were fit to a 1:1 Langmuir binding model, determining association rate (*k*_a_), dissociation rate (*k*_d_), maximum response (R_max_), and the dissociation constant (K_D_) for the interaction.

### Chemicals for cellular BRET

BRET substrate (Nano-Glo®) was purchased from Promega. DMEM, trypsin (0.05%), FCS, and HBSS were purchased from Thermo Fisher Scientific. If not stated elsewhere, all other reagents were purchased from Sigma-Aldrich.

### Plasmids for cellular BRET

Plasmids encoding masGRK3ct-Nluc, Venus(1–155)-Gγ2, and Venus(155-239)-Gβ1 have been described previously^56^ and were kindly provided by Kyrill Martemyanov. Plasmids containing muscarinic acetylcholine receptor 3 (M3) or PTX-S1 cDNA in a pCAGGS backbone were a kind gift from Asuka Inoue.

### Cell lines/Cell culture for cellular BRET

CRISPR/Cas9 generation of HEK293 Δ7 cells was reported elsewhere^36^. Cells were cultured in DMEM supplemented with FCS (10% V/V), penicillin (100 U/mL), and streptomycin (100 µg/mL) and grown at 37 °C and 5% CO_2_.

### BRET assays

Cells were transfected 24h before the experiments using Polyethyleneimine (PEI, 1mg/mL). Transfection was performed according to the manufacturer’s instructions (DNA to PEI solution ratio 1:3). For activation and inhibition testing of G11 heterotrimers, approximately 2.8 x 10^6^ cells were seeded into 10 cm dishes and immediately transfected with plasmids containing the following cDNA (amounts per dish): M3 receptor (1.5 µg), masGRK3ct-Nluc (0.4 µg), wildtype or mutant Venus(156-239)-Gβ1 (0.4 µg), (1-155)-Gγ2 (0.4 µg), wildtype or mutant Gα11 (0.8 µg), PTX-S1 (0.4 µg). Empty pcDNA3.1 vector was added to a total DNA amount of 5 µg per condition. Cells were washed with PBS, detached with trypsin, and centrifuged at 130 x g for 3 minutes. After washing with PBS, cells were centrifuged again and resuspended in BRET buffer (HBSS supplemented with 20 mM HEPES). Approximately 80,000 cells per well were seeded into white 96-well flat bottom plates (Corning, Corning, USA). For carbachol dose-response curves, cells were mixed with Nano-Glo (final dilution 1:1000) and agonist. For inhibitor dose-response curves, cells were incubated with inhibitor in BRET buffer for 15 minutes. All inhibitor dilutions and the buffer control were equalized to the same DMSO content. Afterward, cells were mixed with Nano-Glo (final dilution 1:1000) and agonist. After approximately one minute of incubation, luminescence (475 nm ± 30 nm) and fluorescence (535 nm ± 30 nm) were measured using the BMG PHERAstar® FSX plate reader for 0.48 s once per minute for 5 minutes at 28 °C. The BRET ratio was calculated by dividing fluorescence emission values by luminescence emission values. BRET signals in dose-response curves are means of the fourth and fifth (minute) measurement shown as an increase in BRET ratio over buffer. For measuring heterotrimer stabilization, approximately 350,000 cells per well were seeded into 6-well plates and were transfected 24 h later with plasmids containing the following cDNA (amounts per dish): masGRK3ct-Nluc (0.025 µg), wildtype or mutant Venus(156-239)-Gβ1 (0.2 µg), (1-155)-Gγ2 (0.2 µg), wildtype or mutant Gα11 (1 µg), PTX-S1 (0.05 µg). Empty pcDNA3.1 vector was added to a total DNA amount of 2 µg per well. Cells were washed with PBS, detached by gentle scraping, centrifuged at 500 x g for 5 minutes, and resuspended in BRET buffer (HBSS supplemented with 20 mM HEPES). Approximately 20,000 cells per well were seeded into white 96-well flat bottom plates (Corning, Corning, USA) and mixed with Nano-Glo to a final concentration of 1:1000. After five minutes of incubation, luminescence (475 nm ± 30 nm) and fluorescence (535 nm ± 30 nm) were measured using the BMG PHERAstar® FSX plate reader for 2.88 s every 15 s for 20 minutes at 28 °C. Inhibitor dilutions or DMSO-adjusted BRET buffer were added manually after 120 s of baseline read. The BRET ratio was calculated by dividing fluorescence emission values by luminescence emission values. BRET signals in kinetic measurements are shown as buffer-corrected increase in BRET ratio over baseline by subtracting the mean of the baseline-read and buffer values corresponding to each data point.

## REFERENCES

1. Milligan G, Kostenis E. Heterotrimeric G-proteins: a short history. Br J Pharmacol 147 **Suppl 1**, S46–55 (2006).

2. Oldham WM, Hamm HE. Heterotrimeric G protein activation by G-protein-coupled receptors. Nat Rev Mol Cell Biol 9, 60–71 (2008).

3. Johnston CA, Siderovski DP. Receptor-mediated activation of heterotrimeric G-proteins: current structural insights. Mol Pharmacol 72, 219–230 (2007).

4. Dai SA, et al. State-selective modulation of heterotrimeric Galphas signaling with macrocyclic peptides. Cell 185, 3950–3965 e3925 (2022).

5. Hermes C, Konig GM, Crusemann M. The chromodepsins – chemistry, biology and biosynthesis of a selective Gq inhibitor natural product family. Nat Prod Rep 38, 2276–2292 (2021).

6. Fujioka M, Koda S, Morimoto Y, Biemann K. Structure of Fr900359, a Cyclic Depsipeptide from Ardisia-Crenata Sims. J Org Chem 53, 2820–2825 (1988).

7. Nishimura A, et al. Structural basis for the specific inhibition of heterotrimeric Gq protein by a small molecule. Proc Natl Acad Sci U S A 107, 13666–13671 (2010).

8. Schrage R, et al. The experimental power of FR900359 to study Gq-regulated biological processes. Nat Commun 6, 10156 (2015).

9. Hermes C, et al. Thioesterase-mediated side chain transesterification generates potent Gq signaling inhibitor FR900359. Nat Commun 12, 144 (2021).

10. Annala S, et al. Direct targeting of Galphaq and Galpha11 oncoproteins in cancer cells. Sci Signal 12, (2019).

11. Onken MD, et al. Targeting nucleotide exchange to inhibit constitutively active G protein alpha subunits in cancer cells. Sci Signal 11, (2018).

12. Lapadula D, et al. Effects of Oncogenic Galpha(q) and Galpha(11) Inhibition by FR900359 in Uveal Melanoma. Mol Cancer Res 17, 963–973 (2019).

13. Klepac K, et al. The Gq signalling pathway inhibits brown and beige adipose tissue. Nat Commun 7, 10895 (2016).

14. Matthey M, et al. Targeted inhibition of G(q) signaling induces airway relaxation in mouse models of asthma. Sci Transl Med 9, (2017).

15. Schlegel JG, et al. Macrocyclic Gq Protein Inhibitors FR900359 and/or YM-254890-Fit for Translation? ACS Pharmacol Transl Sci 4, 888–897 (2021).

16. Kostenis E, Pfeil EM, Annala S. Heterotrimeric G(q) proteins as therapeutic targets? J Biol Chem 295, 5206–5215 (2020).

17. Malfacini D, et al. Rational design of a heterotrimeric G protein alpha subunit with artificial inhibitor sensitivity. J Biol Chem 294, 5747–5758 (2019).

18. Patt J, et al. An experimental strategy to probe Gq contribution to signal transduction in living cells. J Biol Chem 296, 100472 (2021).

19. Boesgaard MW, et al. Delineation of molecular determinants for FR900359 inhibition of Gq/11 unlocks inhibition of Galphas. J Biol Chem 295, 13850–13861 (2020).

20. Voss JH, et al. Structure-affinity and structure-residence time relationships of macrocyclic Galpha(q) protein inhibitors. iScience 26, 106492 (2023).

21. Pistorius D, et al. Promoter-Driven Overexpression in Chromobacterium vaccinii Facilitates Access to FR900359 and Yields Novel Low Abundance Analogs. Chemistry 28, e202103888 (2022).

22. Reher R, et al. Deciphering Specificity Determinants for FR900359-Derived Gq alpha Inhibitors Based on Computational and Structure-Activity Studies. ChemMedChem 13, 1634–1643 (2018).

23. Zhang H, Nielsen AL, Stromgaard K. Recent achievements in developing selective G(q) inhibitors. Med Res Rev 40, 135–157 (2020).

24. Randolph CE, et al. Enhanced membrane binding of oncogenic G protein alphaqQ209L confers resistance to inhibitor YM-254890. J Biol Chem 298, 102538 (2022).

25. Maeda S, Qu Q, Robertson MJ, Skiniotis G, Kobilka BK. Structures of the M1 and M2 muscarinic acetylcholine receptor/G-protein complexes. Science 364, 552–557 (2019).

26. Voss JH, et al. Unraveling binding mechanism and kinetics of macrocyclic Galphaq protein inhibitors. Pharmacol Res 173, 105880 (2021).

27. Pandy-Szekeres G, et al. The G protein database, GproteinDb. Nucleic Acids Res 50, D518–D525 (2022).

28. Coleman DE, Sprang SR. Crystal structures of the G protein Gi alpha 1 complexed with GDP and Mg2+: a crystallographic titration experiment. Biochemistry 37, 14376–14385 (1998).

29. Higashijima T, Ferguson KM, Sternweis PC, Smigel MD, Gilman AG. Effects of Mg2+ and the beta gamma-subunit complex on the interactions of guanine nucleotides with G proteins. J Biol Chem 262, 762–766 (1987).

30. Voss JH, et al. Unraveling binding mechanism and kinetics of macrocyclic G alpha(q) protein inhibitors. Pharmacological Research 173, (2021).

31. Majumdar S, Ramachandran S, Cerione RA. Perturbing the linker regions of the alpha-subunit of transducin: a new class of constitutively active GTP-binding proteins. J Biol Chem 279, 40137–40145 (2004).

32. Heydorn A, et al. Identification of a novel site within G protein alpha subunits important for specificity of receptor-G protein interaction. Mol Pharmacol 66, 250–259 (2004).

33. Singh G, Ramachandran S, Cerione RA. A constitutively active Galpha subunit provides insights into the mechanism of G protein activation. Biochemistry 51, 3232–3240 (2012).

34. Darby JF, et al. Water Networks Can Determine the Affinity of Ligand Binding to Proteins. J Am Chem Soc 141, 15818–15826 (2019).

35. Mahmoud AH, Masters MR, Yang Y, Lill MA. Elucidating the multiple roles of hydration for accurate protein-ligand binding prediction via deep learning. Commun Chem 3, 19 (2020).

36. Hisano Y, et al. Lysolipid receptor cross-talk regulates lymphatic endothelial junctions in lymph nodes. J Exp Med 216, 1582–1598 (2019).

37. Lohmann K, et al. Novel GNB1 mutations disrupt assembly and function of G protein heterotrimers and cause global developmental delay in humans. Hum Mol Genet 26, 1078–1086 (2017).

38. Tietze D, et al. Structural and Dynamical Basis of G Protein Inhibition by YM-254890 and FR900359: An Inhibitor in Action. J Chem Inf Model 59, 4361–4373 (2019).

39. Maeda S, et al. Development of an antibody fragment that stabilizes GPCR/G-protein complexes. Nat Commun 9, 3712 (2018).

40. Hanke W, et al. The Bacterial G(q) Signal Transduction Inhibitor FR900359 Impairs Soil-Associated Nematodes. J Chem Ecol, (2023).

41. Abdulrahman W, et al. A set of baculovirus transfer vectors for screening of affinity tags and parallel expression strategies. Anal Biochem 385, 383–385 (2009).

42. Tall GG, Krumins AM, Gilman AG. Mammalian Ric-8A (synembryn) is a heterotrimeric Galpha protein guanine nucleotide exchange factor. J Biol Chem 278, 8356–8362 (2003).

43. Kawate T, Gouaux E. Fluorescence-detection size-exclusion chromatography for precrystallization screening of integral membrane proteins. Structure 14, 673–681 (2006).

44. Vonrhein C, et al. Data processing and analysis with the autoPROC toolbox. Acta Crystallogr D Biol Crystallogr 67, 293–302 (2011).

45. Kabsch W. Xds. Acta Crystallogr D Biol Crystallogr 66, 125–132 (2010).

46. Evans PR, Murshudov GN. How good are my data and what is the resolution? Acta Crystallogr D Biol Crystallogr 69, 1204–1214 (2013).

47. McCoy AJ, Grosse-Kunstleve RW, Adams PD, Winn MD, Storoni LC, Read RJ. Phaser crystallographic software. J Appl Crystallogr 40, 658–674 (2007).

48. Emsley P, Cowtan K. Coot: model-building tools for molecular graphics. Acta Crystallogr D Biol Crystallogr 60, 2126–2132 (2004).

49. Murshudov GN, et al. REFMAC5 for the refinement of macromolecular crystal structures. Acta Crystallogr D Biol Crystallogr 67, 355–367 (2011).

50. Smart OS, Womack, T. O., Sharff, A., Flensburg, C., Keller, P., Paciorek, W., Vonrhein, C. and Bricogne, G. Grade, version 1.2.20.). Cambridge, United Kingdom, Global Phasing Ltd., https://www.globalphasing.com.2011).

51. Davis IW, et al. MolProbity: all-atom contacts and structure validation for proteins and nucleic acids. Nucleic Acids Res 35, W375–383 (2007).

52. Chen VB, et al. MolProbity: all-atom structure validation for macromolecular crystallography. Acta Crystallogr D Biol Crystallogr 66, 12–21 (2010).

53. Joosten RP, Long F, Murshudov GN, Perrakis A. The PDB_REDO server for macromolecular structure model optimization. IUCrJ 1, 213–220 (2014).

54. Varma N, et al. Crystal structure of jumping spider rhodopsin-1 as a light sensitive GPCR. Proc Natl Acad Sci U S A 116, 14547–14556 (2019).

55. Kartal O, Andres F, Lai MP, Nehme R, Cottier K. waveRAPID-A Robust Assay for High-Throughput Kinetic Screens with the Creoptix WAVEsystem. SLAS Discov 26, 995–1003 (2021).

56. Masuho I, Ostrovskaya O, Kramer GM, Jones CD, Xie K, Martemyanov KA. Distinct profiles of functional discrimination among G proteins determine the actions of G protein-coupled receptors. Sci Signal 8, ra123 (2015).

