## Supplementary Data for "Cyclic peptide inhibitors stabilize Gq/11 heterotrimers"

#### Supplementary Figures and Tables

##### Supplementary Figure 1

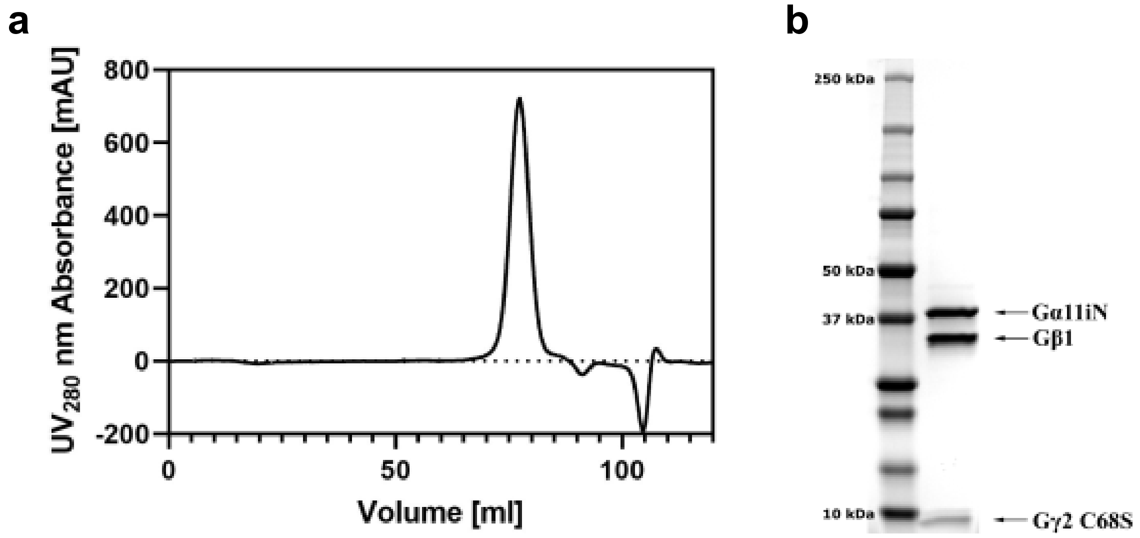

**Supplementary Figure 1: G11iN$_3$-S purification. a)** Final gel filtration chromatogram from a Hiload Superdex 200 pg 160/600 column showing a single, symmetrical, monodisperse peak. **b)** SDS-PAGE analysis (4-20% BioRad MiniProtean TGX gel) of the G11iN$_3$-S trimer prior to crystallization. The sample is highly pure, and the subunits are present in a stoichiometric ratio.

**Supplementary Figure 2**

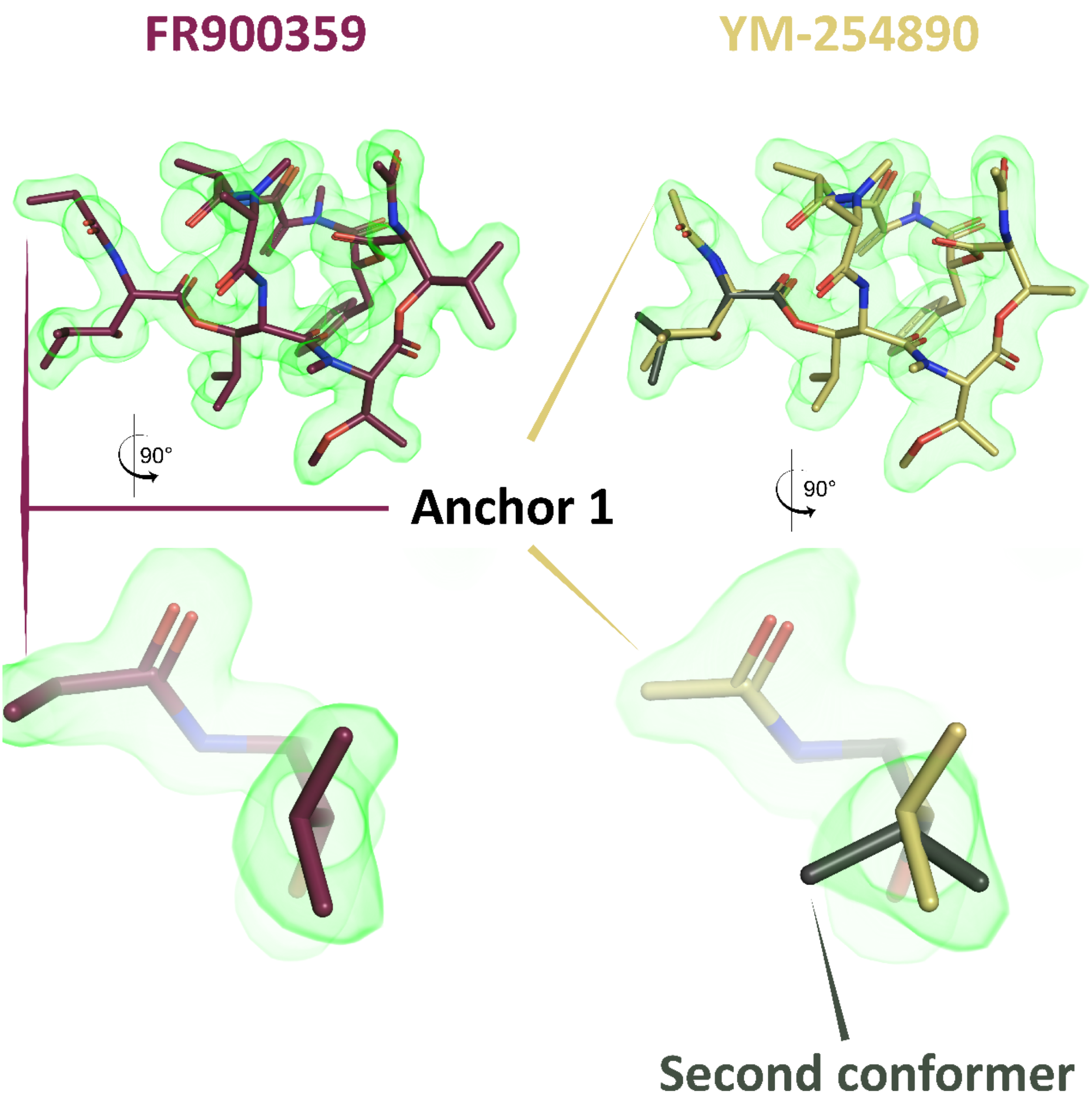

**Supplementary Figure 2: Comparison of the experimental electron densities** of our FR and YM structures with a focus on the second conformer of the β-hydroxyleucine near anchor 1 observed in YM (residue ID: HL2). The simulated annealing OMIT $mF_o - DF_c$ maps are displayed as green volumes contoured at 3.5 σ.

#### Supplementary Figure 3

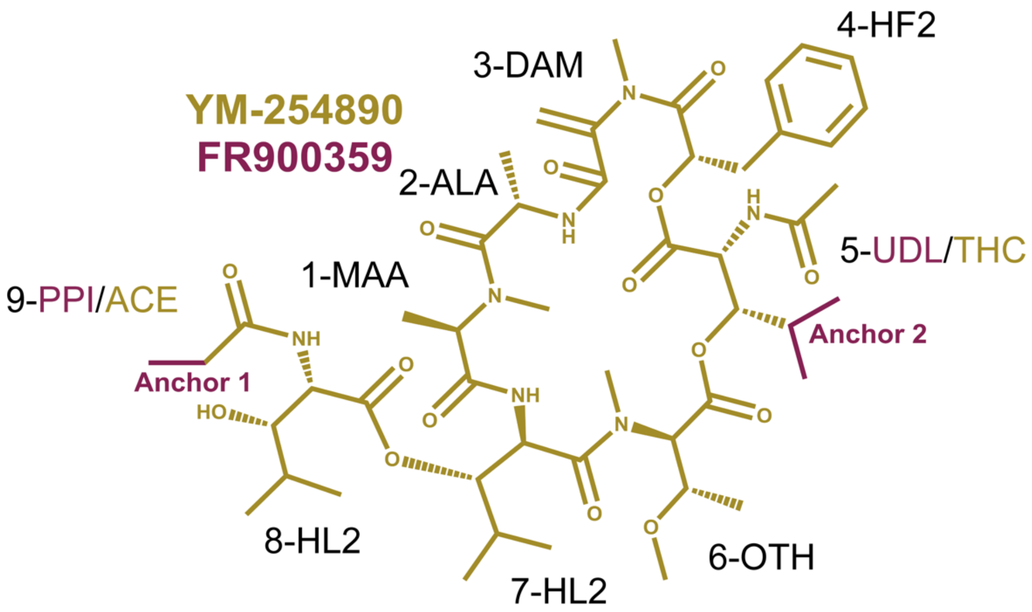

| Residue Number | Three Letter Code | Chemical name | Synonym |
|---|---|---|---|
| 1 | MAA | (2S)-2-(methylamino)propanoic acid | N-methyl-alanine |
| 2 | ALA | (2S)-2-aminopropanoic acid | Alanine |
| 3 | DAM | 2-(methylamino)prop-2-enoic acid | N-methyl-α-β-dehydroalanine |
| 4 | HF2 | (2R)-2-hydroxy-3-phenylpropanoic acid | Phenyllactic acid |
| 5 (FR) | UDL | (2S,3R)-2-acetamido-4-methyl-3-oxidanyl-pentanoic acid | N-acetyl-β-hydroxyleucine |
| 5 (YM) | THC | (2S,3R)-2-acetamido-3-hydroxy-butanoic acid | N-acetylthreonine |
| 6 | OTH | (2S,3R)-3-methoxy-2-(methylamino)butanoic acid | N,O-dimethylthreonine |
| 7 | HL2 | (2S,3R)-2-amino-3-hydroxy-4-methylpentanoic acid | β-hydroxyleucine |
| 8 | HL2 | (2S,3R)-2-amino-3-hydroxy-4-methylpentanoic acid | β-hydroxyleucine |
| 9 (FR) | PPI | Propionic acid | Propionyl |
| 9 (YM) | ACE | Ethanal | Acetyl |

**Supplementary Figure 3. Equivalence between the FR/YM residue identifiers (three-letter code), their chemical names, and common abbreviations.**

**Supplementary Figure 4**

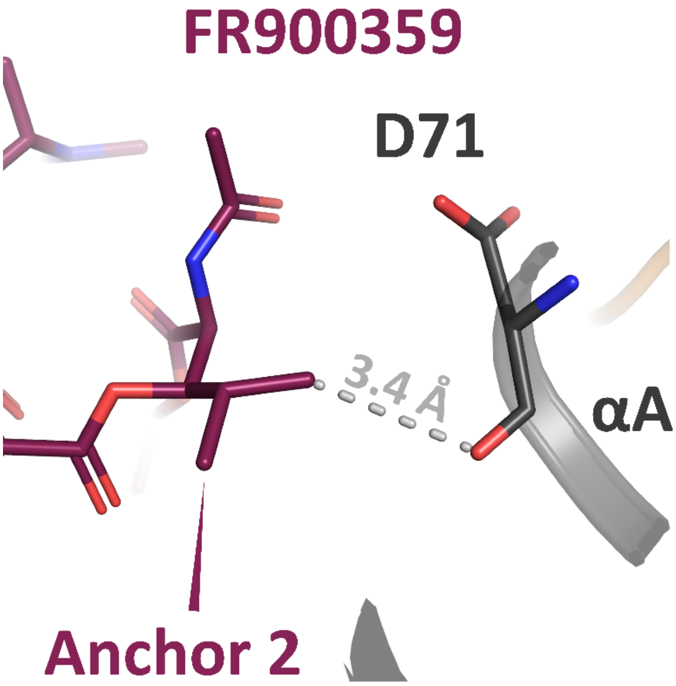

**Supplementary Figure 4: Van-der-Waals interaction of anchor 2 of FR with the backbone carbonyl of Asp71 in the αA helix of the helical domain.** This contact is unique to FR and could be a potential target for FR derivatives that could establish polar interactions in this region.

#### Supplementary Figure 5

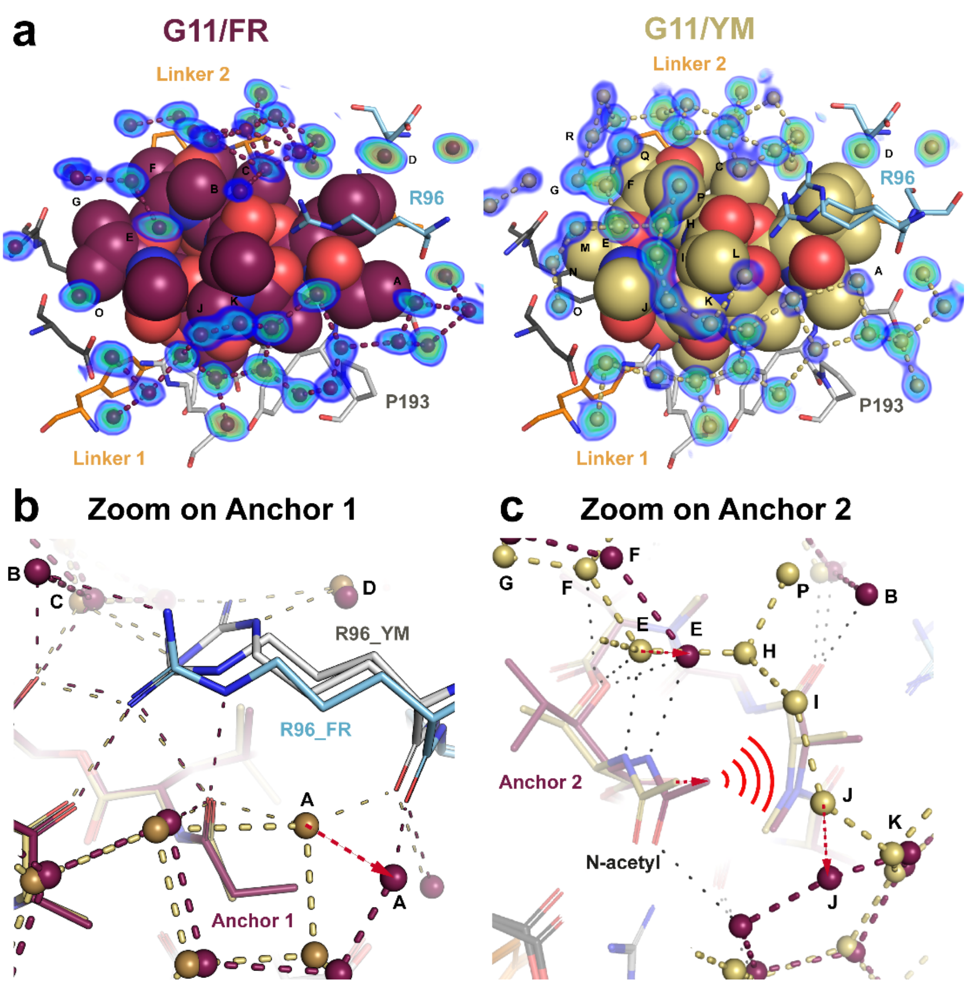

**Supplementary Figure 5: Ligand solvation. a)** Electron densities ($2mF_o - DF_c$) of water molecules around the two ligands are depicted as rainbow-contoured volumes ranging from blue = 1.0 σ to red = 3.5 σ. **b)** Zoom on anchor 1 showing the displacement of water A in the G11/FR structure due to a steric clash with its bulkier anchor 1. **c)** Zoom on anchor 2 showing the desolvation effect of FR in this region triggered by the mild rearrangements of anchor 2 and the nearby FR backbone atoms. In b) and c), water molecules are shown as spheres (FR = red, YM = dark yellow).

#### Supplementary Figure 6

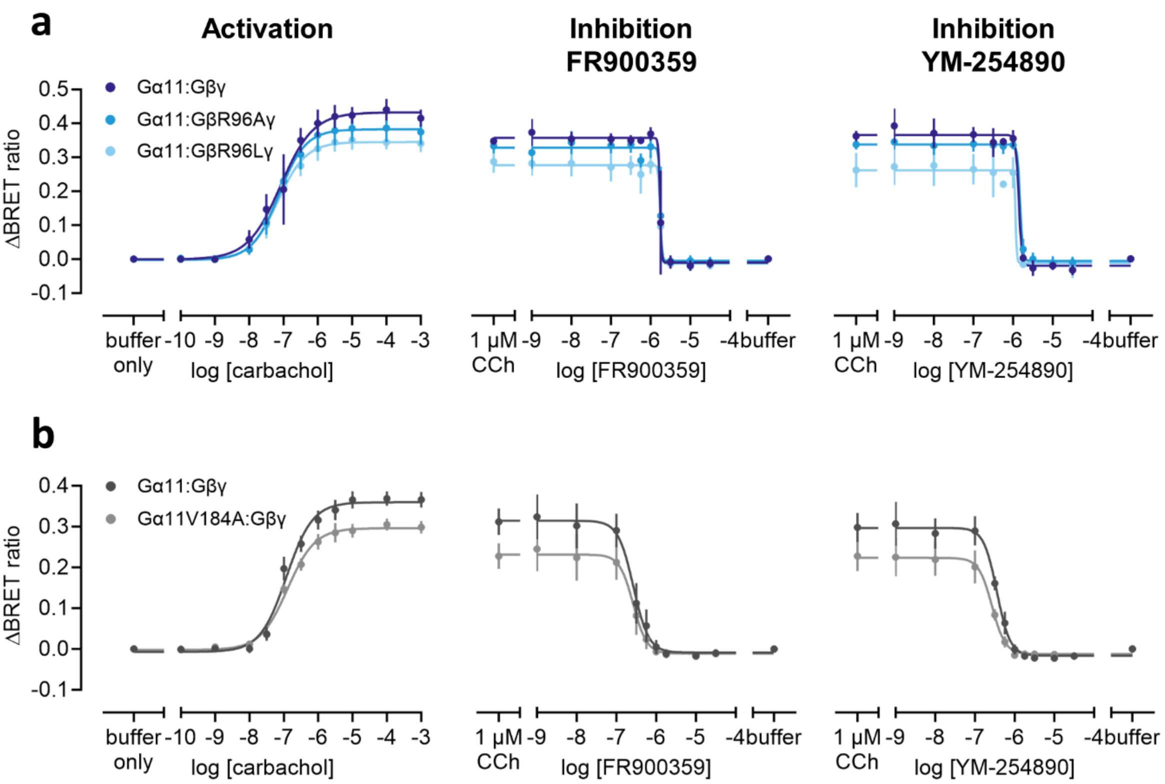

| **c** | **Gα11: Gβγ** | **Gα11: GβR96Aγ** | **Gα11: GβR96Lγ** | **Gα11V184A: Gβγ** |
|---|---|---|---|---|
| $pEC_{50}$ (CCh) | 7.1/6.9 | 7.1 | 7.2 | 6.9 |
| $pIC_{50}$ (FR) | 5.8/6.6 | 5.8 | 5.8 | 6.6 |
| $pIC_{50}$ (YM) | 5.9/6.4 | 5.8 | 5.9 | 6.6 |

**Supplementary Figure 6. Activation and inhibition behavior of the different G11 heterotrimer mutants coincides with wild type. a, b)** Buffer-corrected BRET dose response-curves of carbachol addition (activation via muscarinic acetylcholine receptor M3) and preincubation with FR900359 and YM-254890 (inhibition of carbachol signal induced at submaximal concentration) in HEK293 Δ7 cells transiently transfected with the indicated Gα11 protein and Gβγ split-Venus constructs, Nluc-tagged masGRK3ct, and M3 receptor, as well as PTX-S1 to blunt Gi/o interference. Data points are mean + SE of two to five independent biological replicates performed. **c)** Table of $pEC_{50}$ and $pIC_{50}$ values derived from the dose response-curves in a) and b). BRET: bioluminescence resonance energy transfer, masGRK3ct: membrane-associated GRK3 c-tail; CCh: carbachol.

### Supplementary Figure 7

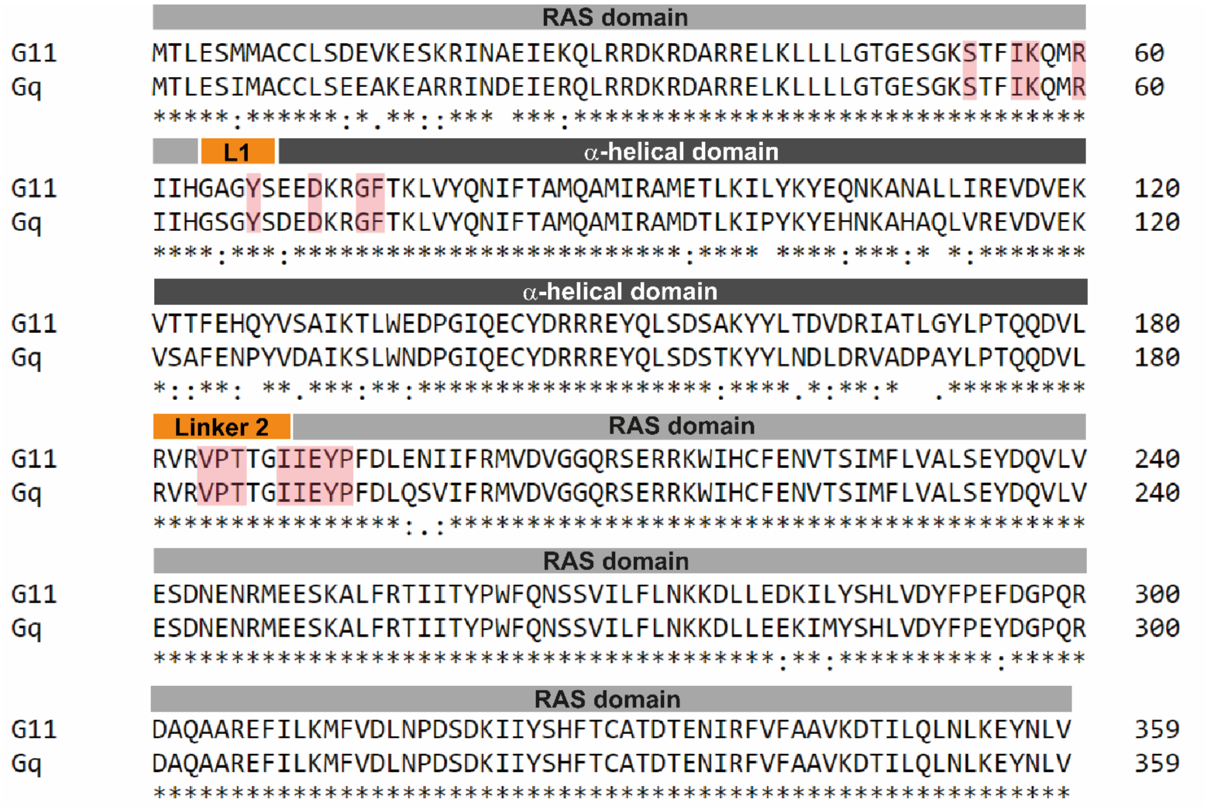

**Supplementary Figure 7: Sequence alignment of human Gα11 vs Gαq.** Identical residues are marked with an asterisk below the sequence. Residues that are interacting with FR are highlighted with red boxes.

### Supplementary Table 1

| **Protein/Ligand** | **G11iN$_3$-S/FR900359** | **G11iN$_3$-S/YM-254890** |
|---|---|---|
| **PBD code** | **8QEH** | **8QEG** |
| *Crystal* | | |
| Space group | P $2_1$ $2_1$ $2_1$ | P $2_1$ $2_1$ $2_1$ |
| Unit cell dimensions (a/b/c in Å) | 72.62/95.80/126.75 | 72.60/102.53/154.00 |
| Unit cell angles (α/β/γ in °) | 90/90/90 | 90/90/90 |
| *Data collection and processing* | | |
| Beamline | SLS-PXIII | SLS-PXI |
| Wavelength (Å) | 1.00 | 1.00 |
| Resolution range (Å)[a] | 126.8-1.43 (1.45-1.43) | 85.35-1.70 (1.73-1.70) |
| Number of unique reflections[a] | 163494 (8016) | 126840 (6244) |
| Completeness (%)[a] | 100.0 (99.9) | 100.0 (100.0) |
| Redundancy[a] | 44.0 (46.7) | 13.7 (14.2) |
| $R_{merge}$(%)[a] | 0.175 (7.774) | 0.059 (1.364) |
| I/σ(I)[a] | 16.5 (0.7) | 22.4 (1.9) |
| $CC_{1/2}$[a] | 1.000 (0.309) | 1.000 (0.796) |
| *Refinement* | | |
| Resolution (Å) | 63.1-1.43 | 59.3-1.70 |
| $R_{work}$ (%) | 0.1401 | 0.1604 |
| $R_{free}$ (%) | 0.1807 | 0.1881 |
| Number of residues | 732 | 726 |
| Number of water molecules | 760 | 804 |
| Average B-factor (Å$^2$) | 28.33 | 38.31 |
| Ramachandran favored (%) | 97.78 | 97.49 |
| Ramachandran allowed (%) | 2.22 | 2.37 |
| Ramachandran outliers (%) | 0.00 | 0.14 |
| RMSD bonds (Å) | 0.01 | 0.01 |
| RMSD angles (°) | 1.53 | 1.69 |

**Supplementary Table 1:** Crystallographic details from data collection, processing, and refinement for the two crystals structures.

[a] Values between brackets are for the highest resolution shell.

#### Supplementary Table 2

| Water | # in G11/FR structure | # in G11/YM structure |
|---|---|---|
| A | 237 | 505 |
| B | 872 | |
| C | 551 | 588 |
| D | 208 | 67 |
| E | 459 | 378 |
| F | 930 | 399 |
| G | 931 | 673 |
| H | | 398 |
| I | | 410 |
| J | 871 | 413 |
| K | 870 | 767 |
| L | | 553 |
| M | | 507 |
| N | | 377 |
| O | 316 | 192 |
| P | | 506 |
| Q | | 408 |
| R | | 797 |

**Supplementary Table 2**: Water numbering in the PDB files for the two structures shown in Fig. 4 of the main text.
